# Genome-wide regulation of KSHV RNA splicing by viral RNA-binding protein ORF57

**DOI:** 10.1101/2022.01.28.478141

**Authors:** Vladimir Majerciak, Alexei Lobanov, Maggie Cam, Zhi-Ming Zheng

## Abstract

RNA splicing plays an essential role in the expression of eukaryotic genes. We previously showed that KSHV ORF57 is a viral splicing factor promoting viral lytic gene expression. In this report, we compared the splicing profile of viral RNAs in BCBL-1 cells carrying a wild-type (WT) versus the cells containing an ORF57 knock-out (57KO) KSHV genome during viral lytic infection. Our analyses of viral RNA splicing junctions from RNA-seq identified hundreds of novel viral RNA splicing events, including the splicing events spanning large parts of the viral genome and production of vIRF4 circRNAs. We found that the 57KO alters the viral RNA splicing efficiency of affected RNA splicing events. Two most susceptible RNAs to ORF57 splicing regulation are the K15 RNA with multiple exons and introns and the bicistronic RNA encoding both viral thymidylate synthase (ORF70) and membrane-associated E3-ubiquitin ligase (K3). ORF70-K3 RNA bears two introns, of which the first intron is positioned in the ORF70 coding region as an alternative intron and the second intron in the intergenic region as a constitutive intron. In WT cells expressing ORF57, most ORF70-K3 transcripts retain the first intron to maintain an intact ORF70 coding region. In contrast, in the 57KO cells, the first intron is substantially spliced out without affecting the splicing efficiency of the second intron. Using a minigene comprising of ORF70-K3 locus, we further confirmed ORF57 regulation of ORF70-K3 RNA splicing, independently of other viral factors. By monitoring protein expression, we showed that ORF57-mediated retention of the first intron leads to the expression of full-length ORF70 protein. The absence of ORF57 promotes the first intron splicing and expression of K3 protein. Altogether, we conclude that ORF57 regulates alternative splicing of ORF70-K3 bicistronic RNA to control K3-mediated immune evasion and ORF70 participation of viral DNA replication in viral lytic infection.

**Author summary:** Like other herpesviruses, the expression of KSHV genes is in a tightly regulated manner during viral infection. The contribution of RNA splicing in the regulation of KSHV gene expression was extensively investigated in this report. We found that approximately one-third of all viral gene transcripts are spliced, with many undergoing RNA alternative splicing. We identified a subset of viral RNA splicing events susceptible to the regulation of viral RNA-binding protein ORF57. This led to the discovery of ORF57-mediated regulation of the bicistronic ORF70-K3 RNA splicing and production of viral thymidylate synthase ORF70 and viral E3-ubiquitin ligase K3. Such regulation allows timely but mutually exclusive expression of these two viral proteins to execute their unique roles in immune evasion and viral genome replication during the KSHV lytic cycle. Thus, these data provide new insight into KSHV biology. Finally, our study represents a blueprint for RNA splicing studies of other pathogens in infected cells.

## Introduction

Kaposi’s sarcoma-associated herpesvirus (KSHV, or human herpesvirus 8, HHV-8) is considered as an etiological cause of several malignancies, including Kaposi’s sarcoma, primary effusion lymphoma (PEL), and multicentric Castleman’s disease [1–3]. KSHV DNA genome encodes up to a hundred viral genes expressing in two functionally distinct transcriptional modes: latent and lytic, each defined by a specific subset of genes with discreet functions in the viral life cycle [4]. The KSHV transcriptome was the subject of numerous studies ranging from single gene locus mapping to genome-wide studies [5–11]. Collectively, these studies form the basis for our current understanding of viral gene expression and its contribution to KSHV pathogenesis. However, many aspects of the regulation in viral gene expression, including contributing viral and host factors, remain to be understood.

RNA splicing plays an essential role in eukaryotic gene expression and almost all human pre-mRNAs are spliced. RNA splicing has been observed in transcripts of numerous viruses, including herpesviruses [12, 13]. However, the number of split genes varies among individual herpesvirus family members, from only a few in alpha-herpesviruses to an estimated ∼30% in KSHV [9, 14]. The level of splicing complexity varies among individual KSHV transcripts, with some exhibiting a splicing pattern resembling cellular constitutive splicing, while others show a highly complex alternative splicing. Functionally, RNA splicing occurs in both coding and untranslated regions (UTR) of KSHV transcripts and regulates viral RNA export and stability and production of different isoforms of proteins or alternative ORFs [15, 16].

Splicing of viral RNAs is mediated by host splicing machinery and thus subjected to regulation by host factors, including components of the cellular spliceosome and host splicing factors. The contribution of a viral factor(s) to RNA splicing was a long time suspected but not systematically studied. We previously showed that KSHV ORF57, a viral RNA-binding protein, acts as a viral splicing factor to promote splicing of several KSHV immediate-early and early transcripts, including ORF50 and K8 [17]. To promote viral splicing, ORF57 interacts with components of host splicing machinery [17]. For example, we showed that ORF57 binds to cellular SRSF3 to relieve its suppressive activity on splicing of K8 suboptimal intron 2 [18], thus preventing the expression of the dominant-negative truncated K8β protein [19]. However, how ORF57 regulates comprehensive RNA splicing in KSHV genome expression remains to be investigated.

The introduction of next-generation sequencing has provided an opportunity to study splicing events across the entire transcriptome. This study aims to provide an unbiased splicing-focused analysis of KSHV transcripts based on RNA-seq analysis of primary effusion lymphoma (PEL) BCBL-1 cells, a patient-derived KSHV-transformed B-cell line [20]. The mapped viral RNA splicing events were analyzed in the context of annotated KSHV genes. To determine the role of ORF57 in KSHV RNA splicing, we compare splicing of viral RNAs in BCBL-1 cells containing a WT KSHV genome with BCBL-1 cells containing an ORF57-null KSHV genome generated by a modified CRISPR/Cas9 knock-out technology [21]. This analysis led to the identification of KSHV splicing events susceptible to ORF57-dependent regulation, including alternative splicing of ORF70-K3 bicistronic transcripts and to uncover a new mode of regulation KSHV gene expression by viral ORF57.

## Results

### Viral gene expression in BCBL-1 single-cell clones with or without KSHV ORF57 expression

To perform a comprehensive genome-wide analysis of KSHV RNA splicing and its regulation by ORF57, we carried out RNA-seq of total RNA isolated from BCBL-1 single-cell clones carrying a wild-type (WT, clone B4) or ORF57 knock-out (57KO, clone #6) KSHV genome 24 h after induction of viral lytic replication with valproic acid (VA24) [21] (**Fig 1A**). Interestingly, for the unknown reason, all selected single-cell WT clones transfected with the empty Cas9 vector express a much lower amount of ORF57 protein when compared to parental BCBL-1 cells but still support KSHV lytic replication (**Fig 1B)**. We selected the WT B4 clone, instead of the WT B8 cells or even the parental BCBL-1 cells, for parallel RNA-seq comparison with the 57KO #6 clone cells because the WT B4 clone cells expressed the lowest amount of ORF57 essential for KSHV lytic replication [21, 22] and thus could give us a similar amount of viral RNA reads for unambiguous RNA splicing analysis of viral RNA transcripts most sensitive to ORF57 regulation. Principal component analysis of differentially expressed genes from three group samples in RNA-seq showed well-separated expression profiles for each group, indicating the high quality of the cDNA library preparation and RNA-sequencing. Each group of four samples exhibited 239 (B4 latent), 233 (B4 lytic), and 236 (57KO #6) million combined high-quality RNA-seq reads (**Fig 1A**). By mapping the sequence libraries to a host GRCh38 (hg38)/BCBL-1 KSHV (GenBank HQ404500) chimeric genome using STAR aligner [23], we found the reduced expression of ORF58 and ORF59 in 57KO cells when compared with WT B4 cells, as expected [24, 25] (**Fig 1C, S1 Table**). Surprisingly, there was only a slight expression decrease (1.3-fold) for PAN RNA from 57KO cells to WT B4 cells (**Fig 1C, S1 Table**), despite that PAN RNA has been identified as a sensitive downstream target of ORF57 [24, 26]. We subsequently verified the RNA-seq results by quantitative RT-qPCR (**Fig 1D**). We showed a ∼73 (68-78)-fold reduction of ORF59 and only 7 (4-10)-fold reduction of abundant PAN RNA in 57KO #6 cells when compared with WT B4 cells. Data suggest the presence of an unknown compensate mechanism for PAN RNA expression in the absence of ORF57 in the course of long-term selection of single-cell clones.

**Fig 1.**
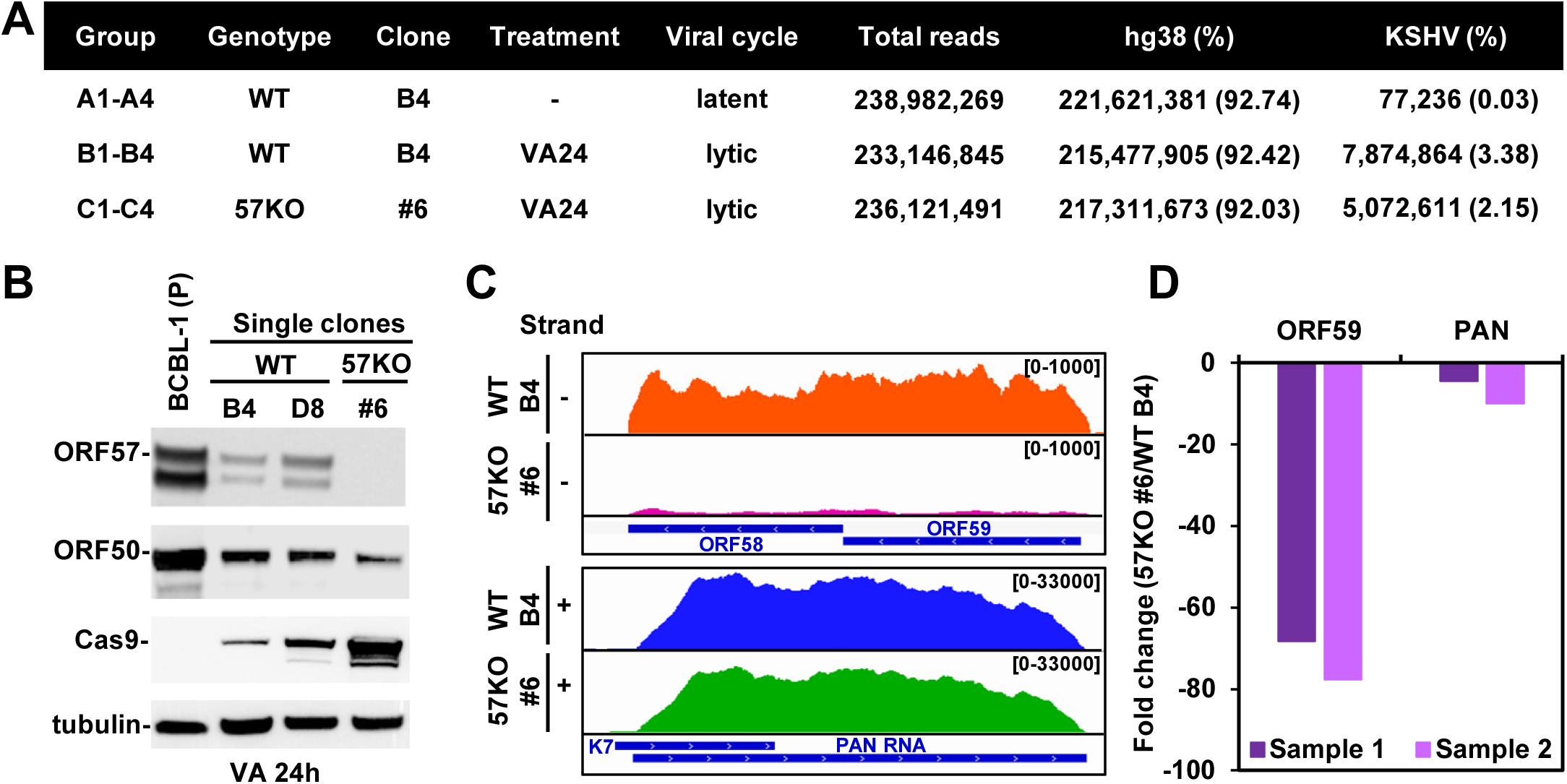
Viral gene expression in BCBL-1 single-cell clones. (A) Samples used for RNA-seq including WT single-cell clone B4 of BCBL-1 cells carrying Cas9 vector lacking gRNAs expression and 57KO single-cell clone #6 carrying a plasmid expressing gRNA2+4 (pABC8) described in previous studies [21]. Total RNA from uninduced clone B4 cells representing WT cells with latent infection (4 samples, A1-4), clone B4 cells treated with 1 mM valproic acid for 24 h (VA24) representing WT cells with the lytic infection (4 samples, B1-4), and clone #6 cells treated with 1 mM valproic acid for 24 h (VA24) representing 57KO cells with the lytic infection (4 samples, C1-4) were subjected to RNA-seq. Total RNA reads uniquely mapped to the hg38/KSHV chimeric genome were counted in each group of four samples. (B) Relative protein expression of ORF57, ORF50, and Cas9 in parental BCBL-1 (P) and BCBL-1 single-cell clones containing a wild-type genome (WT clones B4 and D8) or an ORF57-null genome (57KO clone #6) under 24 h of KSHV lytic infection induced by 1 mM valproic acid (VA 24h). Immunoblot was performed with each protein corresponding specific antibody, with β-tubulin serving as a protein loading control. (C) RNA-seq reads-coverage of KSHV ORF58-59 and PAN RNA in one representative sample each from WT B4 and 57KO #6 cells with KSHV lytic infection induced by 24 h VA treatment. The numbers in the upper right corner represent the read depth. (D) Validation of PAN and ORF59 RNA expression from WT B4 and 57KO #6 cells by RT-qPCR. Total RNA from WT B4 and 57KO #6 cells with KSHV lytic infection described in Panel C was used for the assays.

### Mapping of KSHV splicing junctions from the WT KSHV to ORF57-null KSHV genome in BCBL-1 cells

As our previous study showed high reliability in qualitative and quantitative analysis of genome-wide RNA splicing of mouse papillomavirus (MmuPV-1) [27] by using STAR aligner [23], the splice reads uniquely mapped to the KSHV genome were extracted, pooled together, and used for viral RNA splicing analysis. Because both groups exhibited an almost equal sequence library size, we did not perform any further read counts normalization. In total, we identified 22,978 viral splicing reads derived from 269 unique splicing junctions in the cells with the WT genome and 25,087 viral splicing reads derived from 255 unique splicing junctions in the cells with the 57KO genome (**Fig 2A**). Among those viral splicing junctions, 145 were commonly detected in both WT and 57KO cells. However, many identified splicing junctions were supported with only a few reads and were not consistently detected in all samples within individual groups, indicating they represent rare, low-frequency RNA splicing events (see details in **S2 Table**). Therefore, we next focused only on more prevalent splicing junctions supported by at least 10 or more (≥10) splicing reads. Even though applying this threshold led to ∼5-fold reduction of the number of unique splicing junctions to 57 in the cells with the WT genome and 70 in the cell with the 57KO genome, the splicing reads assigned to these junctions counted for 98% of all identified splicing reads, indicating that these splicing junctions represent the majority of viral RNA splicing events (**Fig 2B**). In addition, most of these junctions (54 in WT and 64 in 57KO) were consistently detected in all four samples within individual groups (**S2 Table**). As expected, the highly prevalent splicing junctions were found in viral RNAs from the WT to 57KO genome, with 50 splicing junctions consistently detected in RNAs expressed from both genomes. The remaining 7 splicing junctions were unique for the WT and 20 for the 57KO genome (**Fig 2B**).

**Fig 2.**
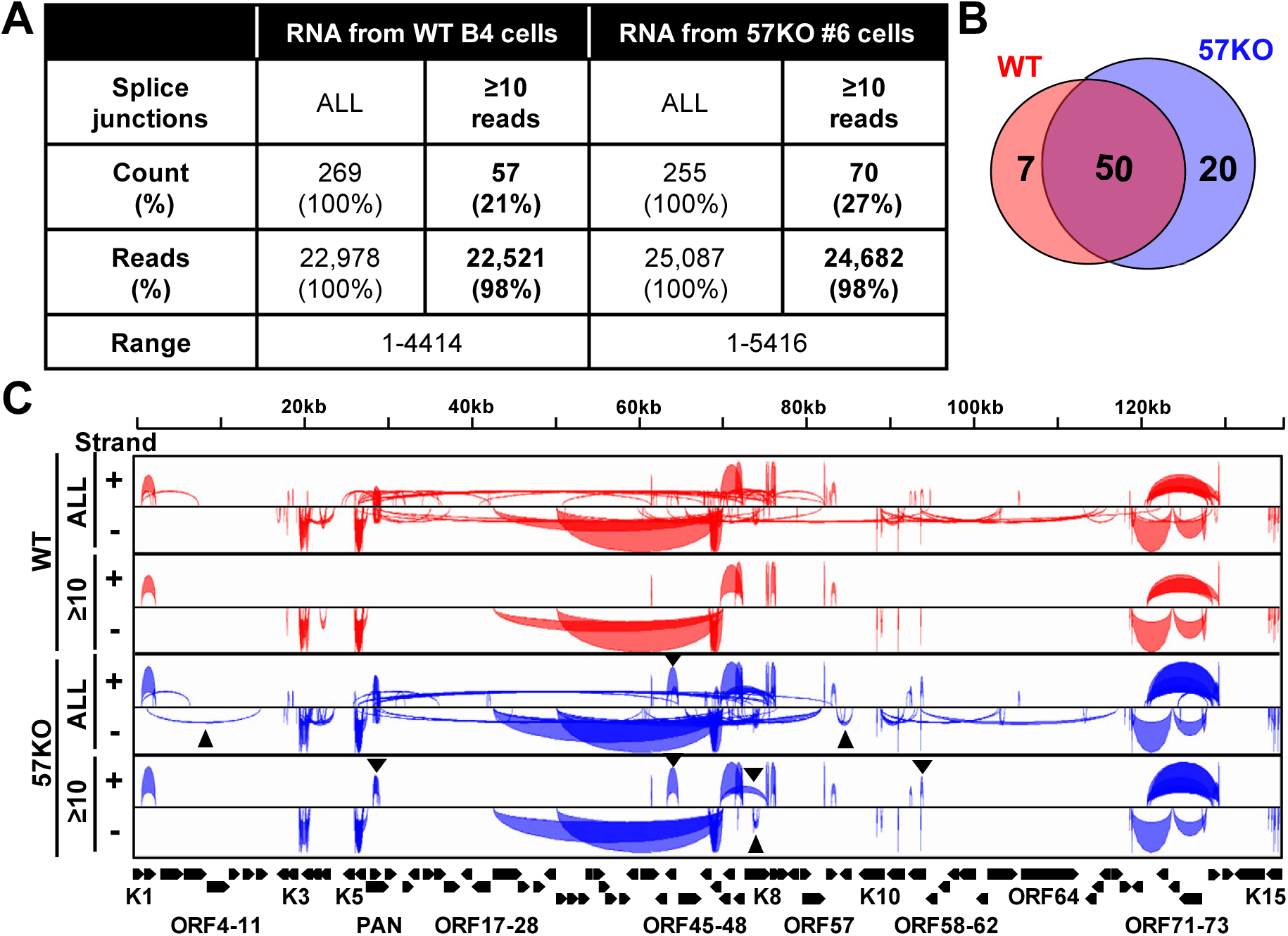
Identification of KSHV splicing junctions by RNA-seq. (A) The number (count) of all detected KSHV RNA splicing junction with the splicing reads (≥10 reads) identified by STAR aligner in WT B4 and 57KO #6 cells (combined 4 samples in each group) with lytic KSHV infection induced by 24 h treatment with 1 mM valproic acid. “Reads” are the number and percentage (%) of all supporting splice reads from all splicing junctions. (B) The Venn diagram shows the overlapped splicing junctions with ≥10 supporting splicing reads between the WT and 57KO genome. (C) The Sashimi graphs generated by IGV showing the viral genomic locations and frequency of RNA splicing events in individual groups. The black triangles indicate the notable RNA splicing differences between the 57KO and WT genome in the cells.

Mapping obtained splicing junctions to the reference KSHV genome revealed a landscape of viral RNA splicing with clusters of split genes being separated by the genome regions lacking splice sites (**Fig 2C**). We also observed several noticeable differences between RNA splicing from the WT to the 57KO genome (**Fig 2C**). To determine the biological relevance of the detected viral splicing events, all highly prevalent splicing junctions were mapped to individual annotated KSHV transcripts with previously mapped or predicted transcriptional start sites (TSS) and polyadenylation cleavage sites [8–10]. The RNA-seq reads-coverage was used to estimate splicing efficiency. Finally, the coding potential of individual spliced RNA isoforms was analyzed to predict the effect of RNA splicing on the viral proteome.

### KSHV RNA splicing profile from the WT genome in BCBL-1 cells

All 57 highly prevalent splicing junctions with ≥10 splicing reads detected in viral RNAs from the WT were named WT-1 to WT-57 in the order from the KSHV genome 5’ to 3’ direction and are summarized in **Table 1**. They mapped to transcripts from 33 annotated KSHV genes (**Fig 3A and 3B**), confirming the previous estimates that ∼30% of KSHV genes are undergoing RNA splicing when expressed [14]. These include almost all previously reported splicing junctions identified either in the single locus or genome-wide studies [9], except the splicing within the ORF29 gene [5]. In addition, several minor alternative splicing junctions previously reported in ORF4 and K8.1 RNAs were detected in this study but did not pass the threshold of ≥10 splicing reads [28, 29]. In contrast, we identified numerous novel splicing junctions which were not previously reported. These include new splicing junctions in viral transcripts of ALT RNA, ORF70-K3, K5, ORF45-48, K10, and a major latent locus, indicating a higher complexity of RNA splicing of these loci than previously proposed (**Fig 3A and 3B**). We also identified several novel splicing events in the genes that were not found to be spliced in previous studies. These include splicing junctions in the ORF2 and K4.1-4.2 loci (**Fig 3B**). Despite their relatively low prevalence, the RT-PCR results support the newly detected introns in the KSHV WT genome in BCBL-1 cells (**S1 Fig**). Other examples of novel splicing junctions detected in our study are the junctions between K5 encoding membrane-based E3-ubiquitin ligase (K5) and a viral cytokine K6 (vMIP-I) [30]. Interestingly, we identified that not only splicing junctions could be mapped to individual K5 (WT-23, -24,-25) and K6 (WT-31,-32,-33) transcripts, but also five new splicing junctions spanning both K5/K6 loci (WT-26,-27,-28,-29,-30) (**Fig 4A**). The existence of these splicing junctions suggests the expression of not only monocistronic K5 and K6 transcripts but also K5-K6 bicistronic transcripts, which are further processed by RNA splicing. The alternative splicing in both K5 and K6 monocistronic RNAs occurs in their 5’ UTRs and is mediated by several alternative 5’ ss and single 3’ ss right upstream of their corresponding coding regions (**Fig 4A**). Interestingly, several upstream ORFs (uORF) expression was detected in both 5’ UTRs [9]. Thus, the observed RNA splicing may regulate K5 and K6 expression by removing these uORFs to efficiently translate K5 and K6 proteins. The splicing of bicistronic RNA occurs between a 5’ ss upstream of K6 ORF and a 3’ ss upstream of K5. This RNA splicing removes the entire K6 code region without affecting the K5 coding capacity. Therefore, the spliced K6^K5 mRNA under control by the K6 promoter encodes only K5 protein (**Fig 4A**). The existence of these RNA splicing events was confirmed by RT-PCR (**Fig 4B**). Together these data reveal a comprehensive profile of KSHV genome-wide RNA splicing events during KSHV lytic infection.

**Fig 3.**
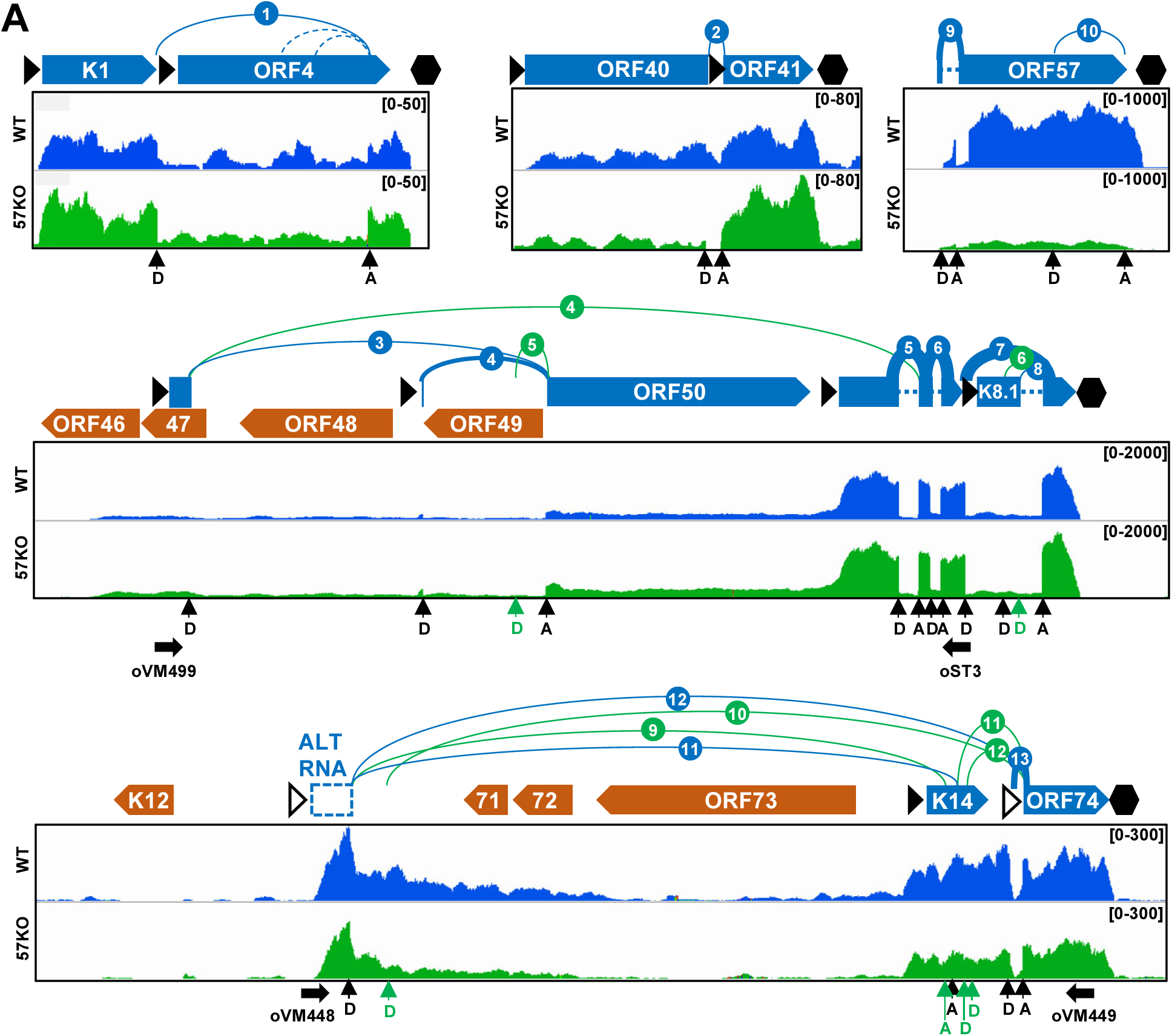
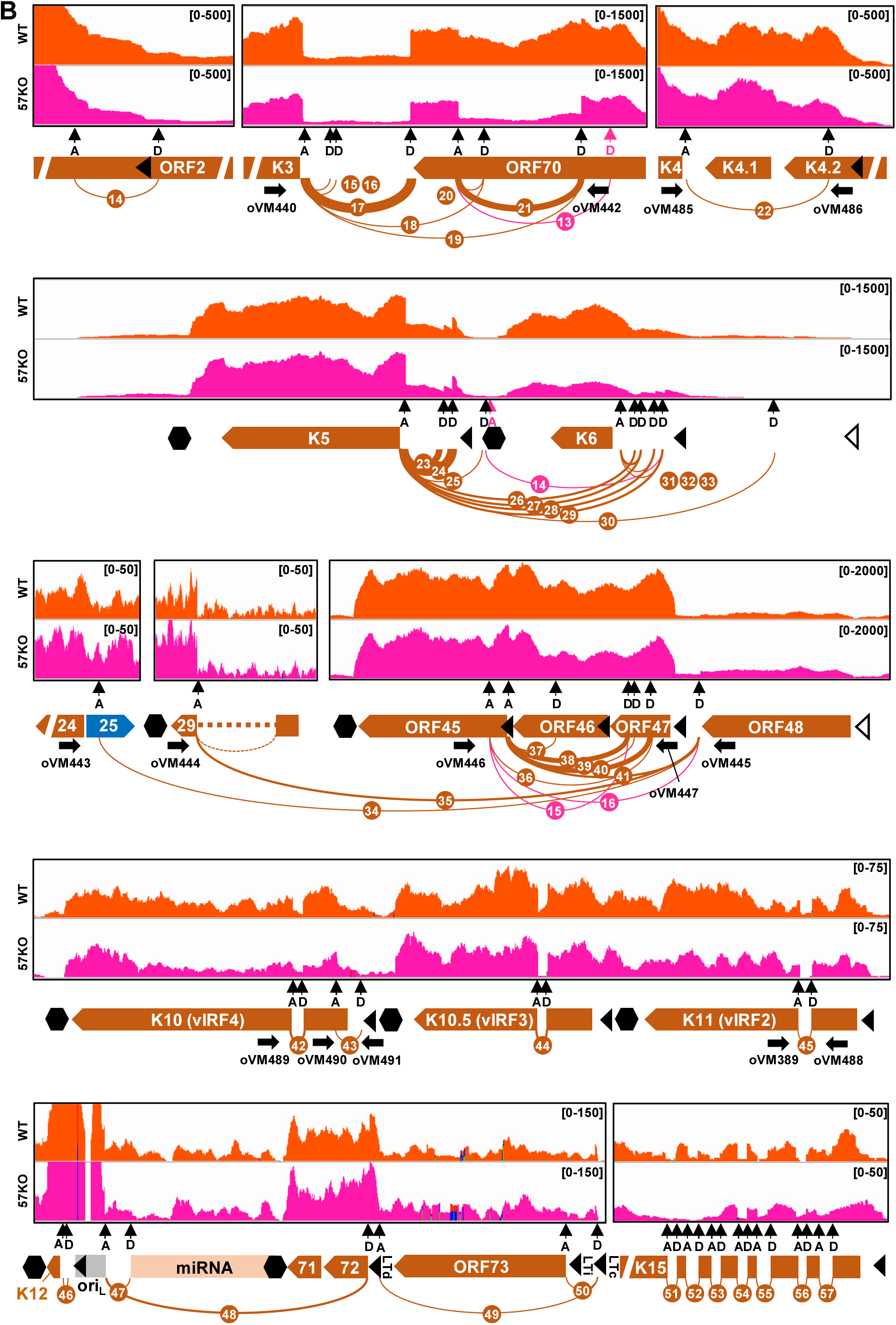
Mapping of KSHV RNA splicing junctions. (A and B) The diagrams depicting all splicing junctions (colored arches) with ≥10 supporting splice reads mapped to viral transcripts expressed from the plus (A) or minus (B) strand of the KSHV genome, with the splicing events being numbered in the order from the genome 5′ to 3′ direction. The blue and orange arches represent splicing junctions detected from the WT genome and green and pink arches from the 57KO genome in the cells (see **Tables 1 and 3**). The arches thickness represents a relative abundance of detected splicing junction reads. The RNA-seq reads-coverage from one representative sample from the WT and 57KO genome in the cells with lytic infection is shown below (A) or above (B) with the reads-coverage depth shown in the upper right corner. The annotated KSHV ORFs are marked by full arrows, blue for ORFs encoded from the plus or orange for ORFs from the minus DNA strand. The dashed boxes are predicted non-coding exons. The black triangles represent mapped (full) or predicted (empty) transcriptional start sites [9, 10]. The black hexagons mark the mapped polyadenylation cleavage sites [8]. The 5′ splice donor (D) and 3′ splice acceptor (A) sites are marked with vertical arrows. Horizontal arrows mark the positions of primer sets used in RT-PCR. LTc - constitutive latent promoter, LTi - inducible latent promoter, LTd - distal latent promoter, oriL - lytic origin of replication.

**Fig 4.**
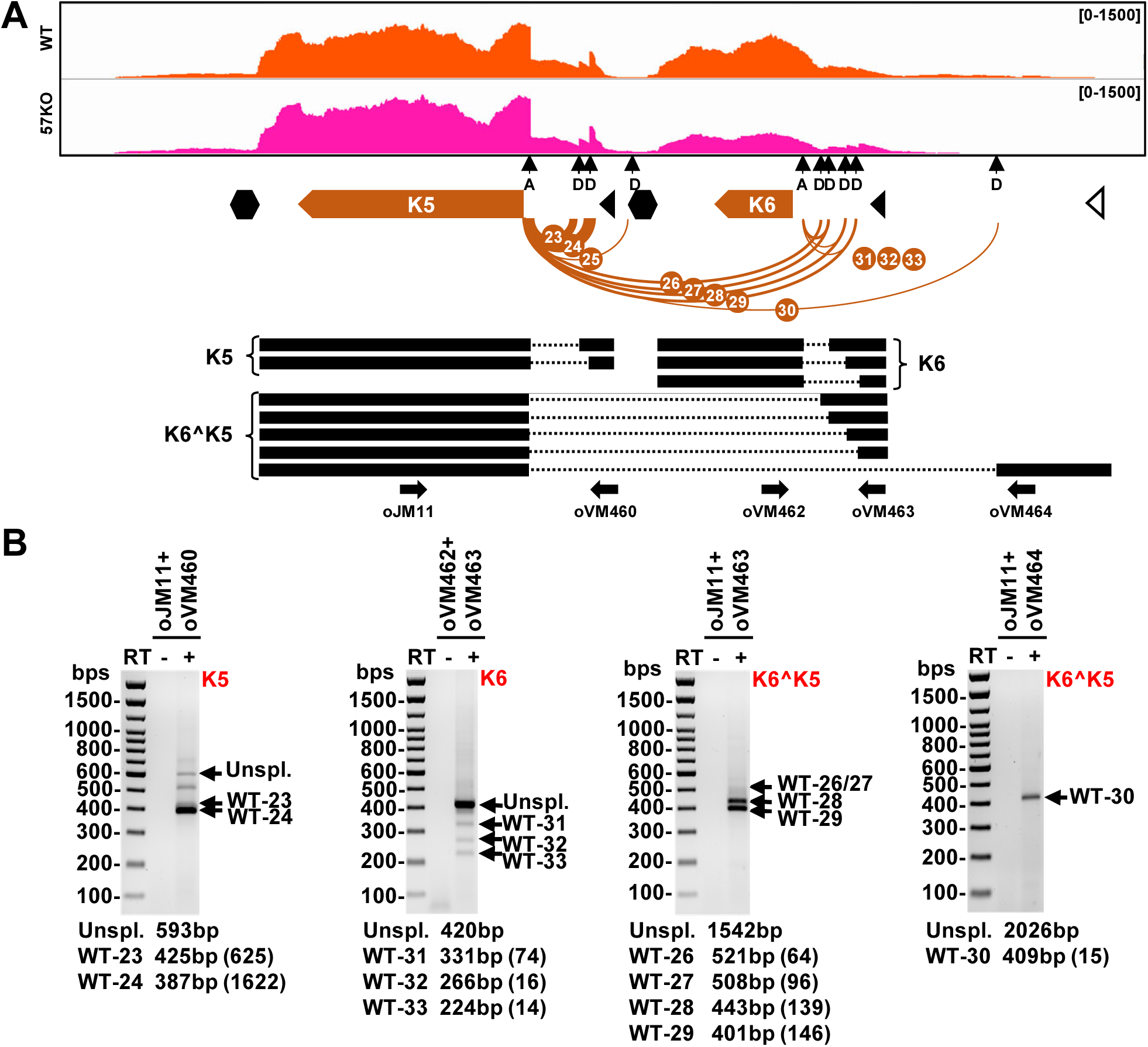
Alternative RNA splicing detected and verified from the K5/K6 locus. (A) The splicing junctions mapped to K5 and K6 locus in the WT genome. Below is the predicted structure of K5, K6, and K5^K6 transcripts with thick black lines representing the exons and dotted lines the introns. The arrows represent the primers used in RT-PCR shown in (B). The predicted sizes of amplificons from unspliced and spliced transcripts are displayed under each gel electrograph. All spliced products amplified were gel purified and confirmed by sequencing. The number of supporting splice reads from the WT genome in the cells with lytic KSHV induction are in the parentheses.

**Table 1:** Splicing junctions detected from the WT genome in BCBL-1 cells supported with ≥10 splice reads. The splicing junctions supported with ≥10 splice reads in BCBL-1 cells with wild type (WT) genome were named WT-1 to -57 from 5′ to 3′ direction and assigned to annotated KSHV genes. The fold change (FC) represents splicing changes in 57KO versus WT cells during viral lytic replication.

| Splicing junction | Intron |  |  | Strand | Reads |  | Locus | FC 57KO/WT |
| --- | --- | --- | --- | --- | --- | --- | --- | --- |
|  | Start (5'ss) | End (3'ss) | Size |  | WT | 57KO |  |  |
| WT-1 | 982 | 2624 | 1643 | + | 28 | 42 | K1/ORF4 | 1.50 |
| WT-2 | 61180 | 61306 | 127 | + | 28 | 27 | ORF40-41 | -1.04 |
| WT-3 | 69365 | 72093 | 2729 | + | 46 | 72 | ORF50 | 1.57 |
| WT-4 | 71136 | 72093 | 958 | + | 467 | 649 | ORF50 | 1.39 |
| WT-5 | 74846 | 74992 | 147 | + | 4414 | 4860 | ORF50/K8 | 1.10 |
| WT-6 | 75086 | 75166 | 81 | + | 3034 | 3796 | ORF50/K8 | 1.25 |
| WT-7 | 75361 | 75954 | 594 | + | 3800 | 5416 | ORF50/K8 | 1.43 |
| WT-8 | 75861 | 75954 | 94 | + | 49 | 42 | ORF50/K8/K8.1 | -1.16 |
| WT-9 | 81641 | 81748 | 108 | + | 1885 | 172 | ORF56/ORF57 | -11.11 |
| WT-10 | 82498 | 83068 | 571 | + | 17 | 15 | ORF56/ORF57 | -1.14 |
| WT-11 | 119872 | 127582 | 7711 | + | 29 | 25 | ALT RNA | -1.16 |
| WT-12 | 119872 | 128428 | 8557 | + | 23 | 99 | ALT RNA | 4.30 |
| WT-13 | 128280 | 128428 | 149 | + | 359 | 223 | K14/ORF74 | -1.61 |
| WT-14 | 18216 | 18095 | 122 | - | 21 | 4 | ORF2 | -5.26 |
| WT-15 | 19695 | 19576 | 120 | - | 23 | 57 | K3/ORF70 | 2.48 |
| WT-16 | 19717 | 19576 | 142 | - | 31 | 36 | K3/ORF70 | 1.16 |
| WT-17 | 20049 | 19576 | 474 | - | 3393 | 2048 | K3/ORF70 | -1.67 |
| WT-18 | 20378 | 19576 | 803 | - | 45 | 7 | K3/ORF70 | -6.25 |
| WT-19 | 20803 | 19576 | 1228 | - | 26 | 38 | K3/ORF70 | 1.46 |
| WT-20 | 20378 | 20264 | 115 | - | 13 | 13 | K3/ORF70 | 1.00 |
| WT-21 | 20803 | 20264 | 540 | - | 691 | 1839 | K3/ORF70 | 2.66 |
| WT-22 | 22789 | 22044 | 746 | - | 12 | 2 | K4.2/K4 | -5.88 |
| WT-23 | 26259 | 26092 | 168 | - | 625 | 511 | K5 | -1.22 |
| WT-24 | 26297 | 26092 | 206 | - | 1622 | 1232 | K5 | -1.32 |
| WT-25 | 26446 | 26092 | 355 | - | 11 | 4 | K5/K6 | -2.78 |
| WT-26 | 27109 | 26092 | 1018 | - | 64 | 65 | K5/K6 | 1.02 |
| WT-27 | 27125 | 26092 | 1034 | - | 96 | 145 | K5/K6 | 1.51 |
| WT-28 | 27190 | 26092 | 1099 | - | 139 | 175 | K5/K6 | 1.26 |
| WT-29 | 27232 | 26092 | 1141 | - | 146 | 215 | K5/K6 | 1.47 |
| WT-30 | 27708 | 26092 | 1617 | - | 15 | 19 | K5/K6 | 1.27 |
| WT-31 | 27125 | 27037 | 89 | - | 74 | 15 | K6 | -5.00 |
| WT-32 | 27190 | 27037 | 154 | - | 16 | 3 | K6 | -5.26 |
| WT-33 | 27232 | 27037 | 196 | - | 14 | 3 | K6 | -4.76 |
| WT-34 | 69683 | 42545 | 27139 | - | 17 | 34 | ORF48/ORF25 | 2.00 |
| WT-35 | 69683 | 50028 | 19656 | - | 64 | 107 | ORF48/ORF29 | 1.67 |
| WT-36 | 69269 | 67969 | 1301 | - | 10 | 36 | ORF47/ORF45 | 3.60 |
| WT-37 | 68505 | 68120 | 386 | - | 64 | 12 | ORF46/ORF45 | -5.26 |
| WT-38 | 69095 | 68120 | 976 | - | 260 | 500 | ORF47/ORF45 | 1.92 |
| WT-39 | 69135 | 68120 | 1016 | - | 13 | 31 | ORF47/ORF45 | 2.38 |
| WT-40 | 69269 | 68120 | 1150 | - | 231 | 833 | ORF47/ORF45 | 3.61 |
| WT-41 | 69683 | 68120 | 1564 | - | 75 | 333 | ORF48/ORF45 | 4.44 |
| WT-42 | 87966 | 87866 | 101 | - | 54 | 25 | K10 (vIRF4) | -2.17 |
| WT-43 | 88557 | 88322 | 236 | - | 24 | 15 | K10 (vIRF4) | -1.59 |
| WT-44 | 90462 | 90369 | 94 | - | 67 | 80 | K10.5 (vIRF3) | 1.19 |
| WT-45 | 93162 | 93042 | 121 | - | 37 | 47 | K11 (vIRF2) | 1.27 |
| WT-46 | 117853 | 117791 | 63 | - | 13 | 15 | K12 | 1.15 |
| WT-47 | 118144 | 117995 | 150 | - | 36 | 71 | K12 | 1.97 |
| WT-48 | 122789 | 117995 | 4795 | - | 96 | 59 | ORF72 | -1.64 |
| WT-49 | 126872 | 122972 | 3901 | - | 27 | 23 | ORF71/ORF72 | -1.18 |
| WT-50 | 126872 | 126374 | 499 | - | 31 | 17 | ORF73 | -1.82 |
| WT-51 | 134284 | 134200 | 85 | - | 20 | 13 | K15 | -1.54 |
| WT-52 | 134468 | 134387 | 82 | - | 18 | 7 | K15 | -2.56 |
| WT-53 | 134658 | 134577 | 82 | - | 14 | 16 | K15 | 1.14 |
| WT-54 | 134888 | 134812 | 77 | - | 34 | 51 | K15 | 1.50 |
| WT-55 | 135066 | 134979 | 88 | - | 29 | 36 | K15 | 1.24 |
| WT-56 | 135392 | 135310 | 83 | - | 19 | 51 | K15 | 2.68 |
| WT-57 | 135594 | 135486 | 109 | - | 12 | 28 | K15 | 2.33 |

### KSHV RNA splice sites and their usage in viral RNA splicing

Together with the RNA branch point, the splice sites (ss) are the essential *cis*-elements required for intron definition and its removal. The sequence composition of the splice sites varies among introns, resulting in a different strength of individual splice sites and thus differences in RNA splicing efficiency. We found that WT KSHV RNA splicing is mediated by a total of 187 distinct donors 5’ ss and 138 acceptors 3’ ss. The splicing junctions supported by ≥10 splicing junction reads in the WT genome are mediated by splicing between 47 distinct 5’ ss and 35 3’ ss (**Table 2**). The introns within the splicing junctions are all canonical “GU-AG” introns. We found that most of the viral 5’ ss participate in splicing a single intron. Only 9 (19%) of all WT 5’ ss use an alternative 3’ ss, with the 5’ ss at nt 69,683 located in ORF47-48 intergenic region being spliced mostly to three alternative 3’ ss (**Fig 3B, Table 2**). Similarly, only 8 (23%) WT 3’ ss are associated with an alternate 5’ ss usage. A striking exception is that the 3’ ss at nt 26,092 upstream of K5 could accept 8 different 5’ ss for various RNA splicing events (**Fig 3B, Table 2**). Thus, the alternative RNA splicing of KSHV transcripts is mainly driven by alternative 5’ ss and 3’ ss usage.

**Table 2.**
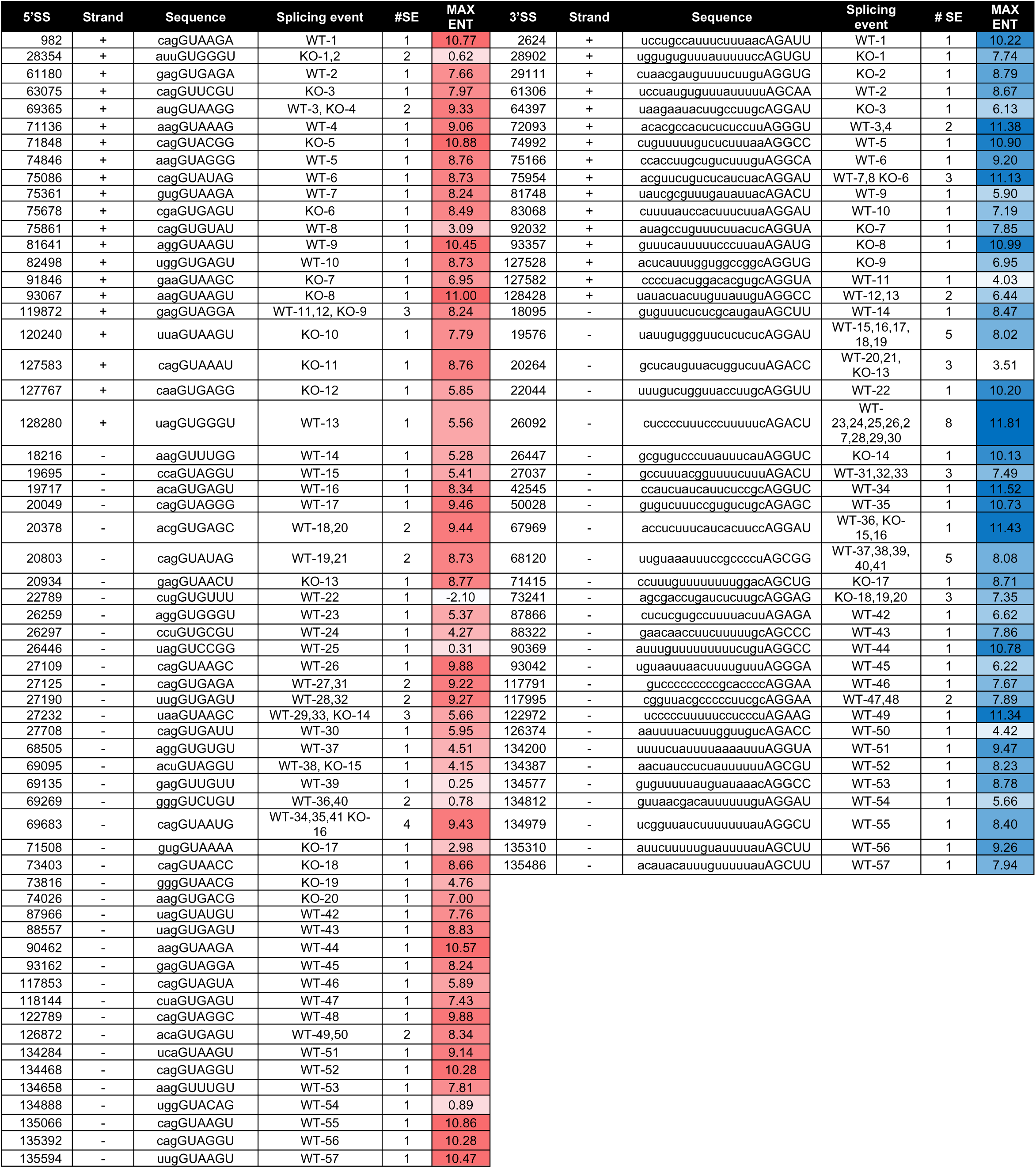
The nucleotide sequences and strength of KSHV 5’ (donor) and 3’ (acceptor) splice sites (ss). The position, strand, sequence, and splicing events (SE) of detected viral splice junctions splice sites supported with ≥10 slice reads. The strength of individual ss was determined by the maximum entropy (MAX:ENT) model [31].

**Table 3:** The 57KO genome-specific splicing junctions supported with ≥10 splice reads. The 57KO-specific splicing junctions supported with ≥10 splice reads named KO-1 to -20 from 5′ to 3′ direction with the assignment to annotated KSHV genes. The fold change (FC) represents splicing changes in 57KO versus WT cells during viral lytic replication.

| Splicing junction | Intron |  |  | Strand | Reads |  | Locus | FC 57KO/WT |
| --- | --- | --- | --- | --- | --- | --- | --- | --- |
|  | Start | End | Size |  | WT | 57KO |  |  |
| KO-1 | 28354 | 28902 | 549 | + | 6 | 30 | PAN | 5.00 |
| KO-2 | 28354 | 29111 | 758 | + | 8 | 26 | PAN | 3.25 |
| KO-3 | 63075 | 64397 | 1323 | + | 2 | 40 | AS-ORF43 | 20.00 |
| KO-4 | 69365 | 74992 | 5628 | + | 6 | 11 | AS-ORF47/K8 | 1.83 |
| KO-5 | 71848 | 72093 | 246 | + | 8 | 43 | ORF50 | 5.38 |
| KO-6 | 75678 | 75954 | 277 | + | 8 | 13 | K8.1 | 1.63 |
| KO-7 | 91846 | 92032 | 187 | + | 3 | 10 | AS-K11 (vIRF-2) | 3.33 |
| KO-8 | 93067 | 93357 | 291 | + | 6 | 26 | AS-K11 (vIRF-2) | 4.33 |
| KO-9 | 119872 | 127528 | 7657 | + | 8 | 11 | ALT RNA | 1.38 |
| KO-10 | 120240 | 128428 | 8189 | + | 7 | 52 | ALT RNA | 7.43 |
| KO-11 | 127583 | 128428 | 846 | + | 0 | 12 | ALT RNA | N/A |
| KO-12 | 127767 | 128428 | 662 | + | 1 | 17 | ALTRNA | 17.00 |
| KO-13 | 20934 | 20264 | 671 | - | 7 | 11 | K3/ORF70 | 1.57 |
| KO-14 | 27232 | 26447 | 786 | - | 7 | 22 | K5/K6 | 3.14 |
| KO-15 | 69095 | 67969 | 1127 | - | 6 | 13 | ORF47/ORF45 | 2.17 |
| KO-16 | 69683 | 67969 | 1715 | - | 3 | 12 | ORF48/ORF45 | 4.00 |
| KO-17 | 71508 | 71415 | 94 | - | 4 | 14 | ORF49 | 3.50 |
| KO-18 | 73403 | 73241 | 163 | - | 4 | 19 | AS-ORF50 | 4.75 |
| KO-19 | 73816 | 73241 | 576 | - | 6 | 10 | AS-ORF50 | 1.67 |
| KO-20 | 74026 | 73241 | 786 | - | 5 | 11 | AS-ORF50 | 2.20 |

To understand the contribution of splice sites to KSHV RNA splicing regulation, we analyze their strength by a maximum entropy (MAX:ENT) model based on sequence composition surrounding 5’ ss and 3’ ss [31]. As shown in **Table 2,** most WT 5’ ss represent the optimal 5’ ss with a median MAX:ENT 8.24, and only six 5’ ss have a suboptimal MAX:ENT below 4.0 (**Table 2**). While most suboptimal intron 5’ ss sites contribute to relatively low-frequency splicing, two of these splice sites mediate RNA splicing to express functional K8.1 (WT-8) and K15 (WT-54) proteins. Their relative weakness could be overcome by their optimal exon size of the upstream exons (K8.1, K15) and presumably the stimulatory effect by splicing adjacent introns (K15) [32]. Like 5’ ss, most WT KSHV 3’ ss are relatively strong, with a median MAX:ENT 8.32 (**Table 2**). Interestingly, the only weak 3’ ss with MAX:ENT under 4.0 was mapped at nt 20,264 within the ORF70 coding region and the splicing of this intron in the ORF70 coding region is subjected to ORF57 regulation (see our data in the late text). Another relatively weak 3’ ss (MAX:ENT 4.42) was found in the LANA 5’ UTR at nt 126,374 and could accept a 5’ ss at nt 126,872 (WT-50) for LANA RNA splicing. Interestingly, the same 5’ ss is also spliced to a much stronger (MAX:ENT 11.34), but distal 3’ ss at nt 122,972 upstream of ORF72/vCyclin (WT-49) [16]. Both RNA splicing events in the polycistronic LANA RNA occur in comparable frequency, indicating that the relative weakness of the proximal 3’ ss in LANA RNA may allow splicing of this 5’ ss to a more distal strong 3’ ss in order to facilitate the expression of all latent proteins from this polycistronic RNA transcribed from the single latent constitutive promoter [15].

### KSHV RNA splicing profile from the 57KO genome in BCBL-1 cells

Similarly, we analyzed the biological significance of viral splicing junctions detected in viral RNAs from the 57KO genome in BCBL-1 cells. As mentioned above, we identified 70 unique splicing junctions with ≥ 10 splicing junction reads from the 57KO genome, of which 50 were also observed from the WT KSHV genome (**Fig 2B**). The remaining 20 splicing junctions were named KO-1 to KO-20 (**Table 3**) and could be split into two groups based on their mapping. The first group represents splicing junctions mapped in viral transcripts with notable splice sites in the WT genome (**Fig 3A and 3B**), but these splice sites could be alternatively spliced to another splice site in the 57KO genome, thus creating a novel splicing junction specific for the 57KO genome. This group of viral RNA transcripts includes the alternatively spliced RNAs from ORF50-K8.1 (KO-4 to -6), ALT RNA (KO-9 and -10), K14-ORF74 (KO-11 and -12), ORF70-K3 (KO-13), K5-K6 (KO-14), and ORF45-48 (KO-15 and -16) (**Fig 3 and S2A**). The second group represents nine novel splicing events not reported or documented in the KSHV intronless RNA transcripts. They were detected from viral lncRNA PAN (KO-1 and -2) and antisense RNA to ORF43 (KO-3), antisense RNA to K11 (KO-7 and -8), ORF49 (KO-17), and antisense RNA to ORF50 (KO-18, -19, -20) (**Fig 5A**). Even though fewer splicing junction reads supported these splicing junctions, they could be readily detected by RT-PCR (**Fig 5B**). It is worth noting that splicing reads associated with most KO splicing junctions could also be found from the WT genome, but with less than 10 reads per junction, except the KO-11 expressed explicitly from the 57KO genome. Together, these data indicate that loss of ORF57 expression led to alternative splicing of KSHV RNAs by activation of cryptic splice sites usage in many intron-containing and/or intronless viral transcripts.

**Fig 5.**
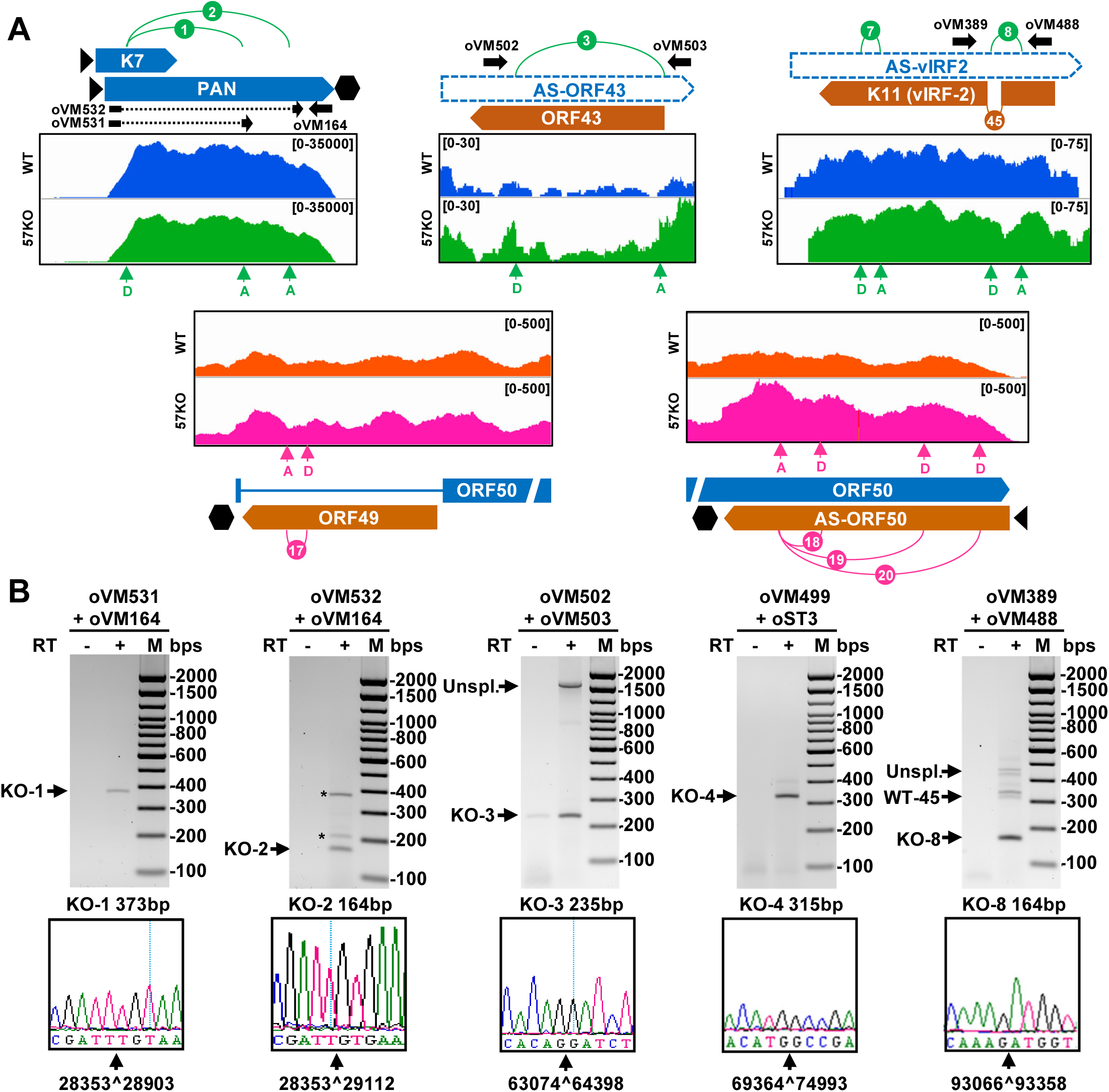
Mapping of 57KO-specific splicing junctions in intronless and antisense viral transcripts. (A) The splicing junctions supported with ≥10 splice reads from the 57KO genome were mapped to intronless viral transcripts. See Fig 3 for a detailed description. (B) RT-PCR validation of selected 57KO-specific RNA splicing events. The sizes of expected amplicons are shown below, together with the chromatographs from Sanger sequencing confirming the mapped splicing junctions.

### Presence of the long-range RNA splicing events crossing over the KSHV genome

Most previously described splicing junctions represent highly prevalent junctions supported by more than 10 splicing junction reads, which were mapped within well-defined transcriptional units (**Fig 3 and 5**). However, we also detected numerous long-range splicing events crossing over the large parts of the virus genome both from WT and 57KO genomes (**Fig 2C**). To better define these splicing events, we first extracted all splicing junction reads covering an intron size ≥10kb and had at least two splicing reads either from one genome or both WT and 57KO genomes (**Fig 6A**). We found a total of 14 such splicing junctions, 7 of them from the WT genome and 10 from the 57KO genome, with 3 found consistently from both genomes in BCBL-1 cells (**Table S2**). A small overlap between WT and 57KO suggests a possible role of ORF57 in regulation in generating these RNAs. Most long-range splicing events occur in low frequency and only two long-range splicing junctions derived from splicing from ORF48 to ORF25 (WT-34) and from ORF48 to the second exon of ORF29 (WT-35) passed ≥10 supporting splicing reads threshold (**Fig 3B, Table 1**). All long-range splicing junctions were generated using canonical splicing sites (**Table S1**). To verify selected long-range splicing events, we performed RT-PCR using various sets of flanking primers on total RNA isolated from both WT and 57KO BCBL-1 cells. Sanger sequencing of the obtained RT-PCR products confirmed the mapped splicing junctions (**Fig 6B**). Together, these data indicate the existence of numerous long-range splicing events in KSHV RNA processing. Although their biological relevance remains to be determined, these long-range spliced RNA isoforms may encode several new proteins (**S3 Fig**) lacking any homology to the known viral proteins worth further investigating in future studies.

**Fig 6.**
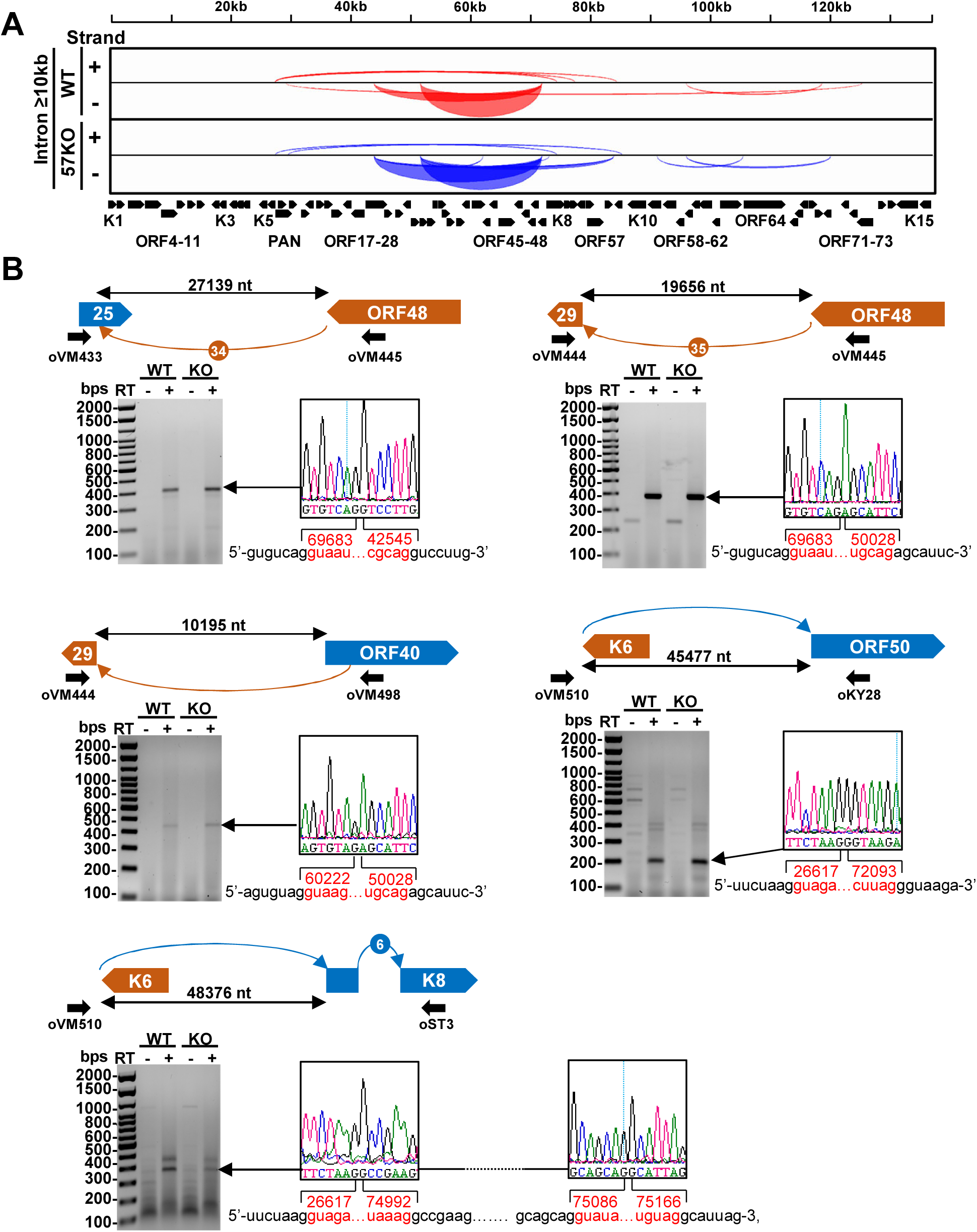
Long-range KSHV RNA splicing identified in BCBL-1 cells with lytic KSHV infection. (A) The Sashimi plot showing all splicing junctions with an intron size ≥10kb supported with at least 2 splicing junction reads. See **S2 Table** for details. (B) Validations of selected long-range RNA splicing events by RT-PCR on total RNA from the WT genome or the 57KO genome in the cells with lytic KSHV replication induced by VA for 24 h. Above is a schematic showing a selected splicing event in orange arches for minus DNA strand-derived RNA transcripts and blue arches for plus DNA strand-derived RNA transcripts, relative genomic locations, predicted size of the intron, and primers used for RT-PCR. Below is the gel electrograph of obtained RT-PCR products. The obtained RT-PCR products were reamplified and confirmed by Sanger sequencing. See sequencing primers in **S6 Table**. The chromatographs on the right show the identified splicing junctions with the exonic sequences marked in black and the intronic sequences spliced out shown in red. The numbers represent 5′ and 3′ ss positions in the KSHV genome (HQ404500).

### Genome-wide KSHV RNA splicing and circular RNA production

RNA splicing usually occurs from the 5’ to 3’ direction and produces a spliced RNA in linear form. However, RNA splicing may occasionally generate a circular RNA (circRNA) by back-splicing when a downstream 5’ ss splices back to the upstream 3’ ss crossing over the exon(s) [33]. Numerous viral circRNAs were reported in KSHV-infected cells; many of them were generated from the RNA transcripts lacking any RNA splicing [34–36]. We subsequently performed viral circRNA prediction from the same RNA-seq library reads using CircRNA Identifier CIRI2 pipeline (https://sourceforge.net/projects/ciri/) [37, 38](**S3 Table**). As shown in **Fig 7A**, we predicted a total of eight unique KSHV viral circRNAs, of which 6 were from the WT and 2 from the 57KO genome. Among 6 circRNAs from the WT genome, 1 mapped to PAN , 4 to antisense PAN (also referred to as K7.3), and one to vIRF4 (K10). Only 2 viral circRNAs were predicted from the 57KO genome, with one mapped to the antisense PAN and the other to ORF57. The supporting back-splicing reads to individual circRNAs were relatively low, ranging from 2 to 9 reads from all samples in each group. All viral circRNAs, except WT-circRNA-1, were predicted in low abundance in BCBL-1 cells by Toptan and colleagues [34]. Given the circRNA being generated by back-splicing from the splice sites used for linear RNA splicing, we next compared all predicted circRNAs in production coordination with those splice sites supported by at least two splicing junction reads identified in our study (**S4 Table**). We found that most viral circular RNAs (**Fig 7A and 7B**) are not supported by the splice sites used for linear RNA splicing, with only WT-circRNA-6 and KO-circRNA-2. As the ORF57 gene is reversely orientated in the 57KO genome [21], the KO-circRNA-2 was genuinely back-spliced by crossing over the exon 2 [39]. We subsequently performed the RT-PCR using several inverse primer pairs on total RNA from the WT and 57KO genome in BCBL-1 cells with lytic induction to verify the predicted viral circular RNAs. Unfortunately, we failed to detect any circRNAs from the PAN region even using 40 amplification cycles and 500 ng of cDNA derived from total RNA extracted either from BCBL-1 cells with a WT or 57KO genome. Even we could not verify any; it is worth noting that the reported PAN circRNAs [34–36] appear mainly next to the ORF57-MRE-I motif we previously identified for PAN RNA stability [26] (**Fig 7B**). We noticed that the predicted PAN circRNAs from Toptan’s report were at different positions from the reported PAN circRNAs in Tagawa’s report (**S4 Fig**) [35]. Ungerleider’s report had no description of PAN circRNAs at all [36]. Subsequently, we analyzed all reported PAN circRNAs with more than 1000 back-splicing junction reads along with our predicted PAN circRNAs for their splice site strength (**S4 Fig**, **S5 Table**). We found none of them giving a reasonable splice site score strong enough to carry out a true RNA splicing reaction (**S4 Fig**). Data suggest that all predicted PAN circRNAs with the current prediction program were unreal.

**Fig 7.**
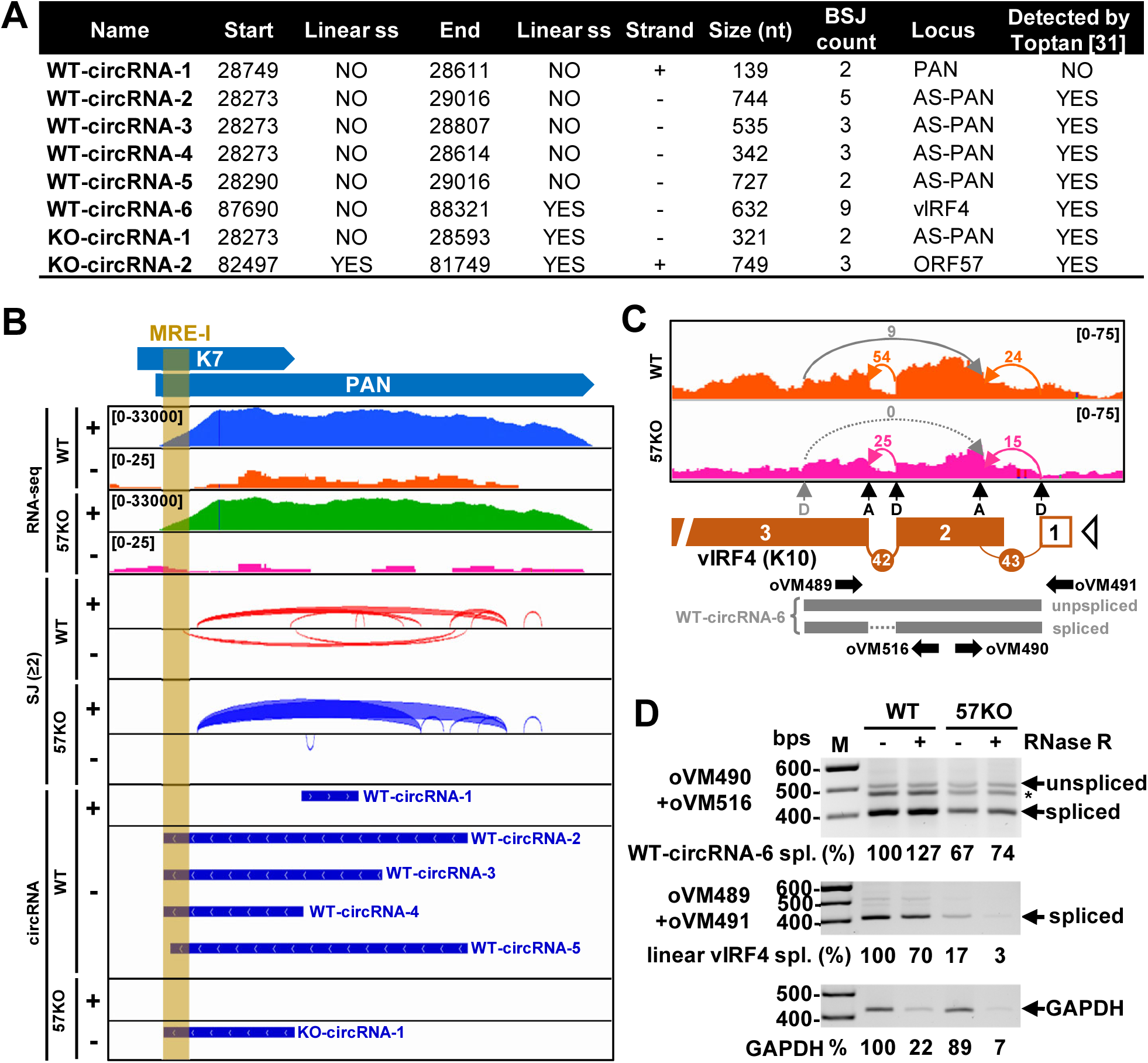
Linear RNA splicing and production of KSHV circRNAs. (A) Identification of KSHV circRNAs from RNA-seq in BCBL-1 cells with a WT or 57KO genome during lytic replication after 24 h valproic acid treatment using a circular RNA prediction software CIRI2 [37, 38]. (B) Distribution and coverage of RNA-seq reads, detected linear RNA splicing junctions (red and blue arches) with minimum 2 splicing junction reads and predicted viral circRNAs (thick blue lines, see panel A for details) in WT and 57KO BCBL-1 cells with lytic replication. MRE-I represents the Mta or ORF57-response element binding ORF57 [26]. (C) The RNA-seq reads mapped to KSHV vIRF4 (K10) locus in WT (orange) and 57KO (pink) cells with reads-depth shown in the upper right corner. The arches show a direction and number of splicing reads detected for linear canonical splicing (orange and pink arches) and backsplicing (grey arches). Below is a diagram of KSHV vIRF4 locus with detected (orange boxes), introns (numbered orange arches) with the corresponding splice sites (D=splice donor and A=splice acceptor). The open triangle represents a predicted transcriptional start site (TSS). Predicted unspliced and spliced forms of WT-circRNA-6 are shown in grey. The primers used to amplify linear and circular vIRF4 RNAs are shown as arrows. (D) The effect of 57KO on the production of vIRF4 linear and circRNAs. The total RNA from WT or 57KO cells isolated 24 h after induction of viral lytic cycle with or without RNase R digestion was used as a template to amplify circular vIRF4 RNA using a primer pair of oVM490 and oVM516 and linear spliced vIRF4 RNA using a primer pair of oVM489 and oVM491 as shown in the panel C. Host GAPDH RNA was amplified as an RNA loading control with the primer pair shown in the **S6 Table**. The percentage of individual RNA was calculated based on band signal density using ethidium bromide-stained agarose gels using ImageLab software (BioRad) with the amount of PCR products obtained from WT RNA without RNase R treatment set as 100%.

Interestingly, the predicted WT-circRNA-6 matches the previously identified circRNA from vIRF4 RNA [34, 36]. We identified two linear RNA splicing events in vIRF4, WT-42, and -43, with WT-42 being about twice more abundant than WT-43 (**Fig 7C, Table 1**). It is worth noting that retention of the WT-43 intron is required to express full-length vIRF4 protein. The 3’ ss of the WT-43 intron in vIRF4 also functions as an acceptor site for the generation of WT-circRNA-6, whereas the donor site used for WT-circRNA-6 production is located in vIRF4 exon 3 and not even used for RNA splicing of linear vIRF4 RNA. As a result, the second intron could be spliced or retained in vIRF4 circRNAs, leading to the generation of two isoforms of circRNA (**Fig 7C**). We observed about two-fold reduction of splicing junction reads in BCBL-1 cells with the 57KO genome when compared with the WT genome and no predicted vIRF4 circRNAs in the cells bearing a 57KO genome. Using different primer sets in RT-PCR and RNase R digestion, we were able to detect and distinguish the linear from circular vIRF4 RNAs and reduced expression of vIRF4 RNA from the 57KO genome in BCBL-1 cells when compared with the WT genome (**Fig 7D**), where the linear vIRF4 RNA in the 57 KO cells appeared ∼80% reduction and accompanied by ∼20% reduction of vIRF4 circRNA. Compared to GAPDH, we also found the linear vIRF4 RNA appeared unexpectedly a little more resistant to RNase R digestion (**Fig 7D**).

### Effect of ORF57 on global KSHV RNA splicing

To fully comprehend the contribution of ORF57 to KSHV RNA splicing regulation, we also compared the number of splicing reads assigned to each splicing junction detected from both WT and 57KO genome (see **Table 1**). Using a minimal 2-fold change cut-off, we identified 21 dysregulated RNA splicing events from the 57KO genome, with 11 being downregulated and 10 being upregulated. As expected, the most downregulated was the splicing of ORF57 first intron (WT-9) due to 57KO [21]. Other downregulated splicing events due to lack of ORF57 expression were mapped to RNA transcripts from ORF2 (WT-14), K3/ORF70 (WT-18), K4.2-K4 (WT-22), K5/K6 (WT-25, -31, -32, and -33), ORF45/46 (WT-37), K10 (WT-42), and K15 (WT-52). The RNA splicing events upregulated in the 57KO cells, ranging from 2.00- to 4.44- fold, were mapped to the transcripts of ALT RNA (WT-12), ORF70/K3 (WT-15 and -21), ORF48/ORF25 (WT-34), ORF47/ORF45 (WT-36, -39, -40), ORF48/ORF45 (WT-41), and K15 (WT-56 and -57). Together, this study discovered that ORF57 has both stimulatory and suppressive effects on KSHV RNA splicing, which could also be confirmed by RT-PCR (**S2 Fig**). Most of the RNA splicing events altered from the 57KO cells were identified from the overlapped viral transcripts bearing a single intron. However, we cannot exclude the possibility that some detected changes might have resulted from the altered expression of affecting genes in the same genome locus. Subsequently, we focused on the altered RNA splicing of KSHV K15 and ORF70-K3, two viral transcripts containing multiple introns and expressed from the genomic loci without overlapping transcription. Thus, we could examine how ORF57 directly affects alternative RNA splicing of these two different transcripts.

### KSHV ORF57 regulates alternative RNA splicing of K15

K15, located at the right end of the long unique region of the KSHV genome, encodes a viral membrane protein with a variable number of transmembrane domains [40, 41]. Through its cytoplasmic C-terminus containing one proline-rich Src homology 3 (SH3) and two SH2-binding sites, K15 interacts with several host factors, such phospholipase Cγ1 (PLC γ1) to activate numerous cellular signaling pathways [42]. K15 is expressed in both latent and lytic infected cells. During latency, K15 promotes the invasiveness of KSHV tumors, KSHV-induced angiogenesis, and the survival advantage of infected cells [43, 44]. By activating PLC γ1, ERK1/2 and AKT1 signaling pathways, K15 promotes reactivation of KSHV from latency and is required for KSHV lytic replication [45, 46]. K15 pre-mRNA harbors eight exons and seven short introns (**Fig 8A**). All internal introns are required to express K15 isoforms with a variable number of transmembrane domains via alternative splicing [40, 41]. We found that the 57KO genome exhibits at least a 2-fold increase of its intron 1 (WT-57) and intron 2 (WT-56) splicing, but a substantial decrease (below 10 splicing junction reads) of the intron 6 (WT-52) splicing (**Table 1**). Using the primer sets shown in **Fig 8B**, the RT-PCR confirmed the increased intron 1 and intron 2 splicing in K15 RNA expressed in the 57KO versus WT BCBL-1 cells (**Fig 8C**). Calculated PSI (percentage spliced-in) for both introns also showed a 2-fold increase of the splicing from the 57KO genome over the WT genome, as shown by RNA-seq analysis (**Table 1**). Consistently, we observed an increased amount of fully spliced K15 RNA expressed from the 57KO genome in the BCBL-1 cells (**Fig 8D**). In contrast, the expression of ORF57 from the WT genome led to an increased level of unspliced K15 RNA, accompanied by decreased level of fully spliced K15 RNA (**Fig 8D**). Retention of the intron 1 or intron 2 from K15 RNA splicing in the presence of ORF57 creates a premature termination codon in these isoforms of K15 RNA, presumably triggering the non-sense mediated RNA decay. We also observed a few K15 RNA alternative introns splicing from the WT to the 57KO genome, but none passed ≥10 reads threshold (**S2 Table**). More specifically, in WT cells, we observed exon 5 skipping and use of an alternative 3’ ss in the exon 8, whereas the cells containing the 57KO genome showed the usage of an alternative 5’ ss in the exon 1 and an alternative 3’ ss in the intron 1. Together, our data indicate that, by regulating K15 splicing, ORF57 might alter the levels of full-length K15 protein and its isoforms during lytic infection.

**Fig 8.**
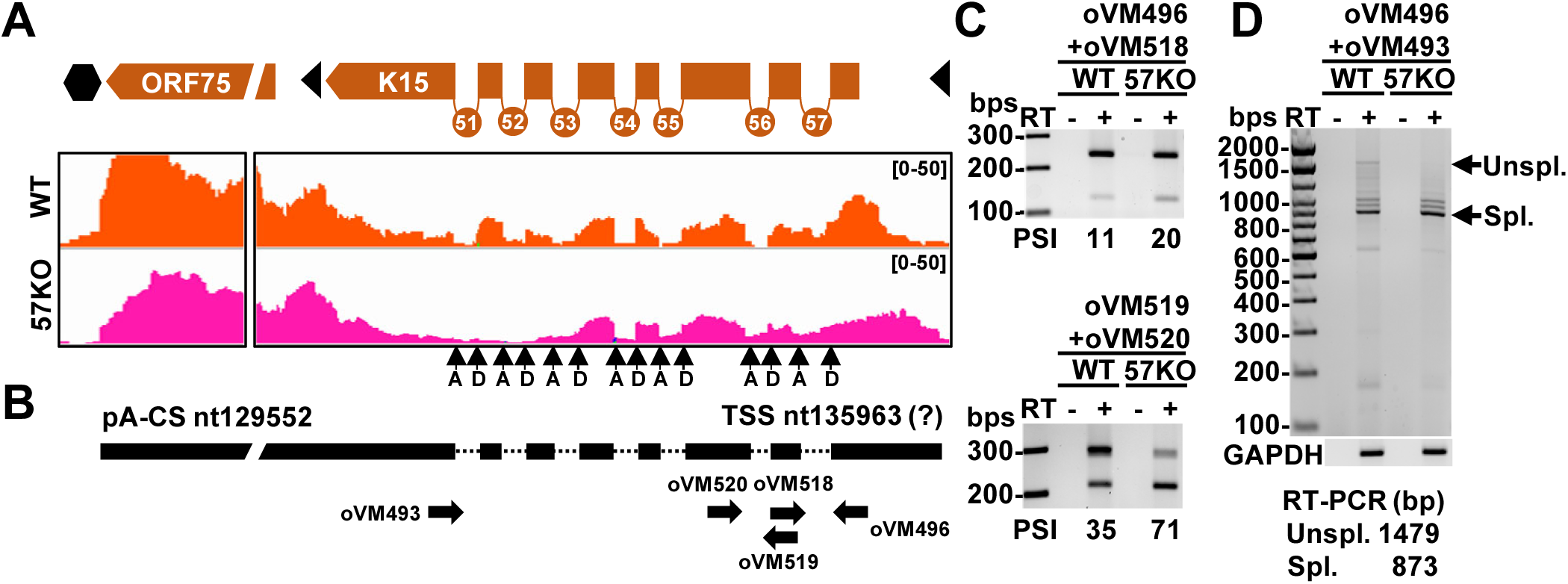
ORF57 regulates K15 alternative splicing. (A) The splicing junctions and RNA-seq coverage of K15 transcripts detected in BCBL-1 cells bearing a WT (orange) or 57KO genome (pink) during KSHV lytic infection. The black arrows mark splicing donor (D) and acceptor (A) sites. (B) The structure of pre-mRNA and fully spliced K15 transcripts with the position of mapped pA cleavage site (CS) and predicted transcriptional start site (TSS). The thick black lines represent the exons and the dotted line the introns. The arrows below show the positions and orientations of the primers used in RT-PCR listed in the **S6 Table**. (C and D) Gel electrographs of RT-PCR products obtained from total RNA from the WT and 57KO genomes in BCBL-1 cells undergoing lytic replication harvested 24 h after VA treatment, with the primer pairs shown in (B). The samples without reverse transcription (- RT) were used as a negative control. The host GAPDH was used as a loading RNA control. PSI, percentage spliced-in; Unspl., unspliced pre-mRNA; Spl., fully spliced mRNA.

### KSHV ORF57 promotes the splicing-dependent expression from K3 to ORF70

ORF70-K3, a bicistronic RNA with complex alternative splicing, is the second viral RNA of which splicing was notably affected from the 57KO genome (**Table 1**). It encodes viral thymidylate synthase ORF70 and viral E3-ubiquitin ligase K3 [47, 48]. We mapped two major introns in ORF70-K3 RNA as reported [6, 49]. The 540 nt-long intron 1 (WT-21) lies within the ORF70 coding region, while the 474 nt-long intron 2 (WT-17) sits entirely into the intergenic region (**Fig 9A**). Similar to previous reports, we also observed low-frequency RNA splicing from the intron 1 donor site to the intron 2 acceptor site (WT-19), resulting in removing a 1228 nt-long intron [6]. Several RNA isoforms generated from this region by alternative RNA splicing were reported in PEL cells with lytic infection [6, 49]. These include an unspliced RNA transcript (isoform A), a single spliced RNA isoform retaining the intron 1 (isoform B), a single spliced isoform retaining the intron 2 (isoform C), a double-spliced isoform (isoform D), and a shortest isoform derived from RNA splicing of the intron 1 donor site to the intron 2 acceptor site (isoform E) (**Fig 9B**).

**Fig 9.**
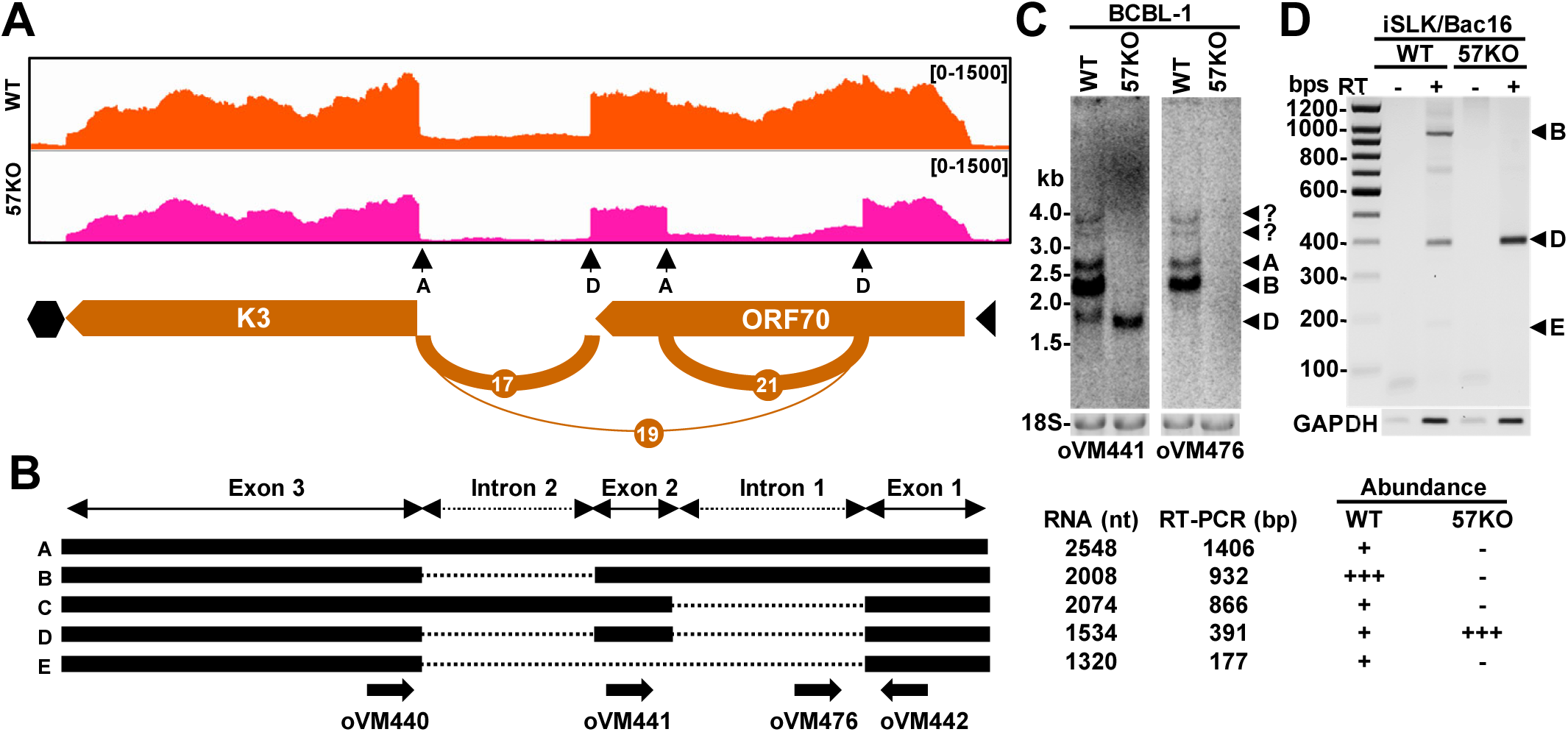
Regulation of ORF70-K3 RNA alternative splicing by ORF57. (A) The most prevalent splicing junctions and RNA-seq coverage of ORF70-K3 transcripts from the WT genome (orange) and the 57KO genome (pink) in BCBL-1 cells during KSHV lytic infection. (B) Diagrams of major ORF70-K3 splicing isoforms. Black boxes represent the exons and dashed lines the introns. The arrows below represent oligos used in Northern blot and RT-PCR (see **S6 Table** for the details). On the right are predicted sizes of all spliced isoforms, sizes of RT-PCR products generated using oVM440 and oVM442 primers, and their relative abundance in the cells with the WT genome or the 57KO genome. (C) Northern blot analysis of ORF70-K3 splicing isoforms expressed from the WT and 57KO genomes in BCBL-1 cells with lytic replication. Total cell RNA was extracted and examined by Northern blot using two separate ^32^P-labeled oligo probes (oVM441 and oVM476) shown in (B). (D) Gel electrograph of RT-PCR products obtained from total RNA from the WT genome or the 57KO genome in iSLK/BAC16 cells during KSHV lytic infection using a primer pair of oVM440 and oVM442 as shown in (B).

Our RNA-seq analysis revealed a significant shift in ORF70-K3 RNA splicing between the WT and 57KO genomes during KSHV lytic infection. In BCBL-1 cells with the WT genome, ORF70-K3 RNA predominantly uses the intron 2 splicing to avoid disruption of ORF70 ORF, with splicing efficiency of the intron 1 being ∼5-fold less than that of the intron 2 (691 intron 1 spliced reads vs 3393 intron 2 spliced reads). On the contrary, in the cells with the 57KO genome, we observed almost equal splicing efficiency for both intron 1 and 2 (1839 intron 1 spliced reads vs 2048 intron 2 spliced reads), indicating a 2.66-fold increase of the intron 1 splicing efficiency but a 1.67-fold decrease of the intron 2 splicing efficiency in the 57KO cells. The splicing between intron 1 and intron 2 showed a minor change between the WT genome (26 reads) and 57KO genome (38 reads) (**Table 1**). We compare the abundance of individual isoforms by Northern Blot on total RNA from the WT and 57KO genome in BCBL-1 cells with lytic replication to confirm these data. First, we use an antisense oligo probe derived from exon 2 (oVM441) to detect all predicted isoforms except isoform E. As shown in **Fig 9C**, three major RNA isoforms were detected from the WT genome, corresponding to the predicted isoforms A (2.5 kb), B (2 kb), C (2 kb), and D (1.5 kb), with the 2 kb transcripts being the most prevalent (**Fig 9B and 9C**). Two minor RNA species between ∼3-4 kb are presumably expressed from the upstream promoters as reported [6, 49]. However, in the cells with the 57KO genome, we detected only a single 1.5 kb RNA corresponding to the double-spliced isoform D (**Fig 9C**). Since this probe could not differentiate the RNA isoform B from the isoform C due to their similar sizes, we reprobed the same membrane with the oligo probe from the intron 1 (oVM476). We observed two major RNA bands in the size of ∼2.0 and 2.5 kb from the WT cells, but not from the 57KO cells, thus confirming their identity as the unspliced isoform A and the single spliced isoform B with retention of the intron 1 in the WT cells (**Fig 9C**). As expected, this probe cannot detect the double spliced isoform D missing both intron 1 and intron 2 (**Fig 9C**).

To investigate if ORF57-mediated regulation of ORF70-K3 RNA splicing is unique to BCBL-1 cells, we took advantage of inducible iSLK cells carrying a KSHV WT or 57KO genomes generated using the same set of gRNAs for 57KO in BCBL-1 cells [21]. Due to the lower viral genome copy number per iSLK cell, the abundance of viral transcripts is significantly lower when compared to BCBL-1 cells. Thereby we were unable to detect ORF70-K3 splice isoforms directly by Northern blot. However, by RT-PCR using a primer pair from the first and last exon of ORF70-K3 bicistronic RNA (see **Fig 9B**), we were able to detect the corresponding RNA isoforms B, D, and E in the cells carrying the WT genome, but only the isoform D in the cells carrying the 57KO genome (**Fig 9D**). These data indicate that ORF57 regulates ORF70-K3 RNA splicing in iSLK/Bac16 cells by inhibiting the intron 1 splicing as observed in BCBL-1 cells.

### KSHV ORF57-mediated ORF70-K3 RNA splicing regulates the production of thymidylate synthase ORF70 and viral E3-ubiquitin ligase K3

Given the bicistronic nature of ORF70-K3 RNA, we next investigated how its RNA splicing regulated by ORF57 affects the expression of individual proteins. As mentioned above, ORF57- sensitive intron 1 lies entirely within the ORF70 coding region thus, only the RNA isoforms A and B retaining the intron 1 could express a 337 aa-long full-length ORF70 protein (ORF70FL, **Fig 10A**). The RNA isoforms C and D lacking intron 1 would express a truncated or spliced ORF70 (ORF70SP) in size of 157 aa residues. ORF70SP lacks the central domain between aa 88 to 269, which bears a catalytic site of thymidylate synthase [50]. The isoform E potentially encodes a further truncated ORF70 (ORF70TR), consisting of only the N-terminal 87 aa residues encoded by the first exon (**Fig 10A**). Interestingly, the K3 coding region for viral E3-ubiquitin ligase remains intact in all spliced RNA isoforms.

**Fig 10.**
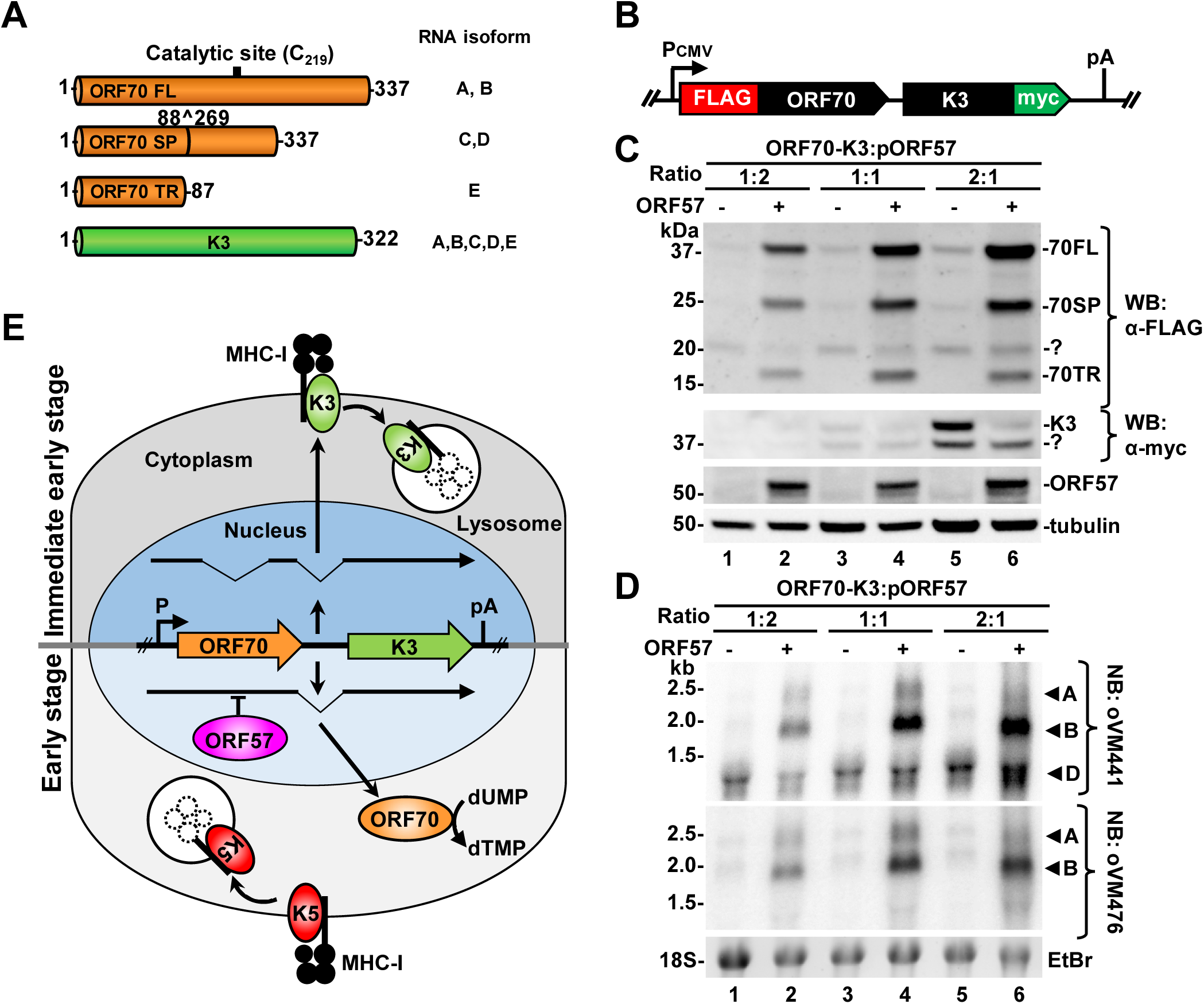
Ectopic ORF57 inhibits alternative splicing of bicistronic ORF70-K3 RNAs to regulate the expression of viral thymidylate synthase ORF70 and viral E3-ubiquitin ligase K3 in HEK293T cells. (A) A number of the amino acid residues of full length (FL), spliced (SP), truncated (TR) ORF70 and K3 proteins expressed from main RNA splicing isoforms are shown on the right (see details in Fig 9B). (B) The diagram of an ORF70-K3 minigene with a FLAG-tag on the ORF70 N-terminus and a c-myc-tag on the K3 C-terminus. The minigene under a CMV IE promoter control contains the viral genomic DNA covering the K3 and ORF70 coding regions and their intergenic region to ensure RNA splicing of this bicistronic RNA transcript. (C and D) Western (C) and Northern blot (D) analyses of HEK293T cells transfected with various ratios of the ORF70-K3 minigene over the ORF57 expression (+) or empty (-) vectors. (C) Expression of ORF70 protein was detected by an anti-FLAG and K3 protein by an anti-c-myc antibody. (D) The changes in ORF70-K3 RNA splicing from the minigene vector in HEK293T cells in the absence (-) or presence (+) of ectopic ORF57 expression were monitored by Northern blot analysis on total cell RNA with two separate oligoprobes shown in Fig 9B. (E) A diagram depicting the role of ORF57 in the regulation of ORF70-K3 RNA alternative splicing to mediate a switch from K3 to ORF70 protein expression during KSHV lytic replication.

Due to the lack of available antibodies against ORF70 and K3, we cannot directly measure the effect of observed regulation of ORF57-K3 alternative RNA splicing by ORF57 on the expression of ORF70 and K3 proteins. Therefore, we constructed a minigene reporter by cloning the ORF70-K3 locus into a mammalian expression vector, allowing us to monitor their expression by introducing two different epitope tags: FLAG-tag on the ORF70 N-terminus and c-myc on the K3 C-terminus (**Fig 10B**). The ORF57-K3 minigene was then transfected into HEK293T cells in various ratios with control empty or an ORF57-expressing vector. Twenty-four after the transfection, the cells were harvested and Western blotted with an anti-FLAG antibody to detect ORF70 and an anti-myc antibody for K3 protein detection. As shown in **Fig 10C**, the minigene in the absence of ORF57 mainly expressed K3 protein and also a small amount of ORF70 protein (lanes 3 and 5). On the contrary, ORF57 co-expression switched from K3 expression to all three isoforms of ORF70 protein, with ORF70FL being the most abundant (**Fig 10C**). The efficiency of the switch was dependent on the ORF57 level and was most prominent at the 2:1 ratio of the minigene plasmid versus the ORF57-expression vector (compare lanes 2, 4, and 6).

We next performed the Northern blot analysis of the corresponding total RNA samples using the same antisense oligoprobes described in **Fig 9B**. As shown in **Fig 10D**, in the absence of ORF57, we found a double-spliced isoform D as a major RNA isoform expressed from the minigene (lanes 1, 3, and 5), which could be confirmed by both exon 2 (oVM441) and the intron 1 (oVM476) probes. In contrast, co-expression of ORF57 was found not only to stabilize all isoform RNAs expressed from the minigenes as reported [22, 26], but also to substantially increase the single spliced isoform RNA B and slightly to increase the unspliced pre-mRNA (isoform A) (lanes 2, 4 and 6). Together, we demonstrated that ORF57-mediated splicing of the minigene-derived ORF70-K3 RNA is responsible for controlling the expression levels of viral thymidylate synthase ORF70 and viral E3-ubiquitin ligase K3, which is independent of other viral factors.

## Discussion

In this study, using RNA-seq analysis combined with bioinformatics and multiple experimental approaches, we performed a comprehensive investigation of viral RNA splicing events associated with KSHV gene expression in BCBL-1 cells carrying a viral WT or a 57KO genome. In addition to detection of all RNA splicing events in the reported gene transcripts [8, 9, 14], we identified hundreds of new splicing events in the expression of both WT and 57KO genomes in the cells with KSHV lytic infection and selectively confirmed most of the newly identified RNA splicing events. By comparing viral RNA splicing events in the expression of the WT over the 57KO genome in BCBL-1 cells, we uncovered the roles of ORF57 in the regulation of KSHV RNA splicing and identified many viral genes as a new target for ORF57-mediated RNA splicing regulation to control KSHV protein expression during lytic infection.

The patient-derived primary effusion (PEL) cell lines represent naturally infected transformed B-cells used as a model for studying KSHV biology, with BCBL-1 cells being one of the most widely used due to the absence of EBV coinfection observed in many other PEL cells [20, 51]. However, the high number of viral genome copies (≥ 100 copies per cell) makes the cell line limited in usage in KSHV genetic studies. We recently developed a modified CRISPR/Cas9 technology allowing the rapid generation of viral ORF57 gene KO from all viral genome copies in selected single-cell clones [21]. These single-cell clones with the 57KO genome allowed us for the first time to study ORF57’s role in viral genome expression during native KSHV infection, thus providing an advantage over the studies conducted in BAC-based recombinant KSHV genome-transfected cell lines [17, 52]. Although ORF57 has been demonstrated to stabilize PAN RNA, we found that the selected 57KO #6 cells appear compensating the expression of genes highly sensitive to post-transcriptional regulation by ORF57, such as PAN of which expression was found just slightly lower than the PAN level in the WT cells during viral lytic infection. The mechanism of such compensation in viral gene expression in the 57KO #6 cells remains to be investigated.

RNA splicing represents an important step in the processing of both eukaryotic and viral pre-mRNAs and is prevalent in the production of numerous KSHV mRNA transcripts [9, 10, 14]. However, the scale, regulation, and biological roles of KSHV RNA splicing remain poorly understood. We identified hundreds of new KSHV RNA splicing junctions, mainly in the expression of viral lytic genes, with 57 highly confident splicing events detected from the WT genome and an additional 20 from the 57KO genome. The newly identified RNA splicing events were found in both split and intronless transcripts (**Fig 3 and 5**). Among ∼30% KSHV split genes undergoing RNA splicing regulation in the virus genome, the majority are KSHV-specific genes (K1-K15) encoding host mimics to tackle host responses during virus infection. These genes presumably originated from the host genome and the high splicing frequency detected in their RNA transcripts may represent an evolutionary memory of their cellular origin. On the contrary, the transcripts from other viral genes being highly conserved in other herpesviruses mostly avoid RNA splicing.

Although the new RNA splicing events discovered in the coding regions of ORF2, K4.2, ORF45-47, ORF70, and K12 may lead to the expression of potentially new protein isoforms (**Fig 3**), other mapped splicing events occur in the untranslated regions (UTR), such as the 5’ and 3’ UTRs or in the intergenic regions (**Fig 3**), thus providing an additional layer of regulation of viral gene expression. For example, we observed extensive alternative RNA splicing from the 5’ UTR of viral K5 and K6 transcripts (**Fig 4**) that harbor several upstream ORFs (µORF) or small ORFs (sORF) [9]. The µORFs and sORFs have been shown to regulate the expression of both host and viral RNAs by stimulating or suppressing the translation of downstream ORFs [53–55]. How the RNA splicing in the 5’ UTRs of viral transcripts contributes to their expression remains to be determined. Additional alternative RNA splicing from K6 5’ UTR to K5 RNA allows K5 expression from a K6 promoter (**Fig 4**) and thus prevents the K6 promoter-derived transcripts from polyadenylation at a pA site mapped downstream of the K6 ORF [8].

KSHV has a common strategy to express a cluster of genes using a gene-specific promoter, but all RNA transcripts transcribed from these different promoters are polyadenylated at a single pA site [8]. The use of this strategy is well documented in the expression of RTA, K8, and K8.1, or ORF48, ORF47, ORF46, and ORF45 genes [56, 57]. In this case, the mapped RNA splicing occurs within transcripts’ 3’ UTRs (**Fig 3**). Here RNA splicing may be required to remove the suppressive and toxic sequences such as the binding site for host miRNAs or RNA-binding proteins, to promote their expression of these polycistronic transcripts [58]. It may also prevent the formation of extended double-stranded RNA structures in a long 3’ UTR recognizable by host RNA sensors which could induce innate immune responses. Interestingly, we recently found that KSHV developed numerous mechanisms to counteract these processes by ORF57-mediated inhibition of P-bodies and stress granules [59, 60].

We have identified a group of splicing junction reads spanning large parts of the viral genome (≥10kbs) across multiple transcriptional units. This long-range RNA splicing happens in both plus-strand RNA transcripts and minus-strand RNA transcripts, predominantly in the same transcription direction but also in the opposite direction of RNA transcripts. The biological relevance of this long-distance RNA splicing remains to be investigated. The coding potential analysis identified several short novel ORFs identified at splicing junctions (**S3 Fig**). Because of their low abundance, the biological relevance of these novel ORF-encoded small proteins also needs to be studied.

Although the presented study focuses on profiling KSHV genome-wide RNA splicing of all viral RNA transcripts from RNA 5’ to 3’ direction (canonical splicing), the multiple reports recently suggested the presence of detectable RNA back-splicing from RNA 3’ to 5’ direction in KSHV transcripts leading to expression of many viral circular RNAs [34–36]. Using a circRNA prediction pipeline CIRI2 [37, 38] to analyze our RNA-seq library reads, we predicted only 8 unique KSHV circRNAs in BCBL-1 cells with lytic replication, six from the WT genome and two from the 57KO genome (**Fig 7A, S3 Table**), mainly from the antisense PAN (or K7.3 )[61]. Despite that both transcripts from PAN RNA locus were not previously considered to be spliced, we observed several rare splicing events in PAN and K7.3 transcripts more often in 57KO cells (**Fig 7B, S1 Table**) but did not find any of these rare splicing events being used for the production of the predicted PAN circRNAs (**Fig 7A**). Experimentally, we could not detect these PAN or K7.3 circRNAs in lytic BCBL-1 cells by RT-PCR using inverse primer pairs as reported [34]. The prediction of K7.3 circRNAs was even more surprising since, in contrast to the highly expressed PAN transcript, expression of this antisense transcript was only minimal in the cells with lytic KSHV infection (**Fig 7B**). The inconsistency among different studies and lack of functional splice sites (**S5 Table**) further suggest that all PAN circRNAs predicted in the recent reports might be the artifacts of RNA-seq library preparation from the erroneously ligated chimeric reads and/or reads analysis software [62].

On the contrary, the back-splicing reads detected from vIRF4 (WT-circRNA-6) locus were real and more comparable to the number of linear RNA splicing junction reads (ratio 1:5 in WT cells) and the overall transcription level (**Fig 7C**). Its 5’ end overlaps with the 3’ acceptor site of vIRF4 first intron (WT-43), spliced to a new 5’ donor site downstream of the second intron (**Fig 7C**). We confirmed the production of vIRF4 circRNAs in WT B4 cells during KSHV lytic infection by RT-PCR using an inverse primer pair next to the back-splice sites. Interestingly, the production of this circRNA was related to ORF57 expression. We found that the reduced expression of vIRF4 circRNAs was correlated with the reduction of linear vIRF4 RNA splicing in 57KO #6 cells (**Fig 7D**). However, the active role(s) of ORF57 in the biogenesis of viral vIRF4 circRNAs remains unknown.

The regulation of KSHV splicing is poorly understood. Viral transcripts are spliced by cellular RNA splicing machinery and thus are subjects to similar regulatory mechanisms as host RNAs, including splice site selections, RNA *cis*-elements in interaction with host splicing factors, etc. [32]. We previously identified viral RNA-binding ORF57 protein as a viral splicing factor promoting the alternative RNA splicing of several early lytic genes, including ORF50 and K8 [17]. A particularly potent effect of ORF57 was observed on the splicing of the suboptimal K8 intron 2 to promote the expression of fully functional K8α [17]. ORF57 regulation of K8 RNA splicing was found to be independent of other viral proteins but required ORF57 binding to host splicing factor SRSF3 to attenuate its suppressive activity on splicing of the K8 intron 2 [18]. Interestingly, we did not observe similar regulation on K8 intron 2 splicing from the 57KO genome in BCBL-1 cells when compared to the WT genome (**Fig 3A, Table 1**). The cause of this discrepancy remains to be understood and might be resulted from the compensated expression of PAN RNA sponging host splicing factors in the 57KO cells. Nevertheless, we found that ORF57 exhibits both stimulatory and suppressive activity on viral RNA splicing and splice site selection (**Table 1 and 3**), a similar dual effect on RNA splicing, which was also observed for ORF57 homolog ICP27 from herpes simplex viruses [63, 64].

The two most remarkable effects of ORF57 on KSHV RNA splicing were identified for RNA processing of monocistronic K15 and bicistronic ORF70-K3 RNA transcripts (**Fig 8 and 9**). K15 gene comprises eight exons and seven introns [14] and encodes a non-structural membrane-bound protein responsible for KSHV-induced angiogenesis and productive infection [42, 46]. We found that the 57KO genome displayed more efficient K15 splicing of the first two introns, both containing a premature termination codon (**Fig 8C**). Their retention, clearly observed from the WT genome in the cells, would reduce the expression of full-length K15 protein. We also observed alternative RNA splicing of K15 RNA, leading to the production of various spliced RNA isoforms as previously reported, but none of these RNA isoforms had enough splicing junction reads over the threshold ≥10 reads. In the WT B4 cells, we observed K15 exon 5 skipping and alternative 3’ ss usage in the K15 exon 8. In the 57KO #6 cells, we found splicing from an alternative 5’ ss in the first exon to a 3’ ss of the intron 3. These data clearly indicate that ORF57 regulates RNA splicing and expression of the K15 gene to fine-tune its activity.

Bicistronic ORF70-K3 RNA transcript encodes viral thymidylate synthase (ORF70) and viral membrane-bound E3-ubiquitin ligase (K3), each being expressed from an individual ORF separated by a 480-nt long intergenic region. In accordance with previous reports, we detected multiple spliced isoforms of ORF70-K3 RNA generated by alternative splicing of two major introns, with a suboptimal intron 1 in the ORF70 coding region and a strong intron 2 within the intergenic region (**Fig 3 and 9**). We found that the intron 1 is alternatively spliced and ORF57 strongly inhibits its splicing. The intron 2 is a constitutive intron not susceptible to ORF57 regulation (**Fig 9**). Consistent with our observation, a similar switch in spliced RNA isoforms of ORF70-K3 RNA expression was reported after treating BCBL-1 cells with cycloheximide, a protein synthesis inhibitor, to suppress the expression of early viral genes, including ORF57 [49]. However, the switch in production of individual ORF70-K3 isoforms in the reported study [49] was wrongly interpreted due to alternative promoter usage.

Functionally, the intron 1 retention in the presence of ORF57 is necessary for the expression of ORF70 during viral lytic infection, but the double spliced RNA isoform D is responsible for K3 protein production. Using an ORF70-K3 minigene, we confirmed that the double spliced RNA isoform D expresses K3 protein, while the intron 1 retention in the ORF70 ORF region mediated by ORF57 led to the expression of ORF70 protein (**Fig 10B-D**). The biological significance of KSHV ORF57-mediated regulation of the expression of ORF70 and K3 is highlighted in **Fig 10E**. In the absence of ORF57, K3 expressed from the double spliced bicistronic ORF70-K3 RNA as a viral immediate-early viral protein functions as an E3-ubiquitin ligase promoting KSHV immune evasion by degrading host MHC-1 proteins to establish productive viral infection [65]. The rising expression of ORF57 in the early stage of virus infection blocks the intron 1 splicing to promote the production of viral thymidylate synthase (ORF70) for robust viral DNA replication by mediating dTMP synthesis [48]. As reported, the expression of structurally and functionally homologous K5 protein can compensate for the loss of K3 expression at later infection [66, 67].

In conclusion, we have identified KSHV ORF57 as a viral splicing factor in the global regulation of KSHV genome expression and RNA splicing. The ORF57-mediated splicing regulation profoundly affects the expression of affected genes as shown in the expression of viral thymidylate synthase ORF70 and viral E3-ubiquitin ligase K3 during the KSHV life cycle, thus providing further insight into KSHV biology.

## Materials and methods

### Cell cultures

KSHV-positive BCBL-1 cells [20] were cultivated in RPMI 1640 medium (Thermo Fisher Scientific). The iSLK cells carrying KSHV BAC16 genomes [67] and HEK293T (ATCC) cells were cultivated in a DMEM medium (Thermo Fisher Scientific). All media were supplemented with 10% fetal bovine serum (FBS, Hyclone, Cytiva) and 1 × Penicillin-Streptomycin-Glutamine (Thermo Fisher Scientific). Hygromycin B (150 μg/ml) and G418 (250 μg/ml) were added to the culture media of iSLK/BAC16 cells to maintain the KSHV genome. The ORF57 knock-outs (57KO) in BCBL-1 and iSLK/Bac16 cells were generated by a modified CRIPS-Cas9 technology as described [21]. The single clones of parental cells transfected with a plasmid expressing Cas9, but lacking gRNA expression were also selected in parallel and used as control wild-type (WT) cells. To reactivate KSHV lytic replication, BCBL-1 cells were treated with 1 mM final concentration of sodium salt of valproic acid (VA, Sigma-Aldrich) for 24 h. KSHV replication in iSLK/Bac16 cells was reactivated by a combination of 1 μg/ml doxycycline (DOX, Clontech) and 1 mM sodium butyrate (Bu, Sigma-Aldrich). RNA samples were collected 24 h after the reactivation. All plasmid transfections were performed by LipoD293 In Vitro DNA transfection reagent (SignaGen Laboratories) as recommended.

### RNA-seq and data analysis

Total RNA from 4 replicates of BCBL-1 WT clone B4 and 57KO clone #6 cells were isolated by TriPure Reagent (Roche) 24 h after induction with VA. The genomic DNA was removed using RNeasy Mini Kit (Qiagen) on-column DNase-treatment. Total RNA-seq libraries were prepared with TruSeq Stranded Total RNA Library Kit and subjected to pair-end sequencing using HiSeq3000/4000 chemistry (2 × 150 nts modality). Reads were trimmed with CutAdapt v.1.18 and mapped to host GRCh38 (hg38)/KSHV (HQ404500) chimeric genome using STAR aligner v.2.7.0f. [23]. The changes of viral gene expression between individual groups were determined by the Limma-Voom package using viral subreads counts mapped to annotated KSHV ORFs. The splice reads mapped to the KSHV genome were extracted to identify unique viral splice junctions. The distribution of splicing junction along the KSHV genome was visualized using Integrative Genomics Viewer (IGV, Broad Institute). The strength of viral splicing sites was calculated using the MaxEntScan::score algorithm [31]. The original sequence libraries were deposited in GEO repository (GSE194239).

### RT-PCR and RT-qPCR

Before RT, the total RNA isolated by TriPure Reagent (Roche) was treated with Turbo DNase (Thermo Fisher Scientific). The specific products were amplified using two-step RT-PCR using random hexamers and primers listed in the **S6 Table**. The reactions without reverse transcriptase were used as the negative controls. The RT-PCR products were separated in agarose gel, purified, and the splicing junctions were verified by Sanger sequencing. For RT-qPCR, the DNase-treated RNAs were converted to cDNAs by SuperScript First-Strand Synthesis System (Thermo Fisher Scientific) using random hexamers as recommended. The resulting cDNAs were used as templates in qPCR reactions containing TaqMan Universal PCR Master Mix (Thermo Fisher Scientific) and custom PrimeTime qPCR assay (IDT) for KSHV PAN and ORF59 RNAs [25, 26]. The cellular GAPDH level detected by the TaqMan assay (Hs02758991_g1, Thermo Fisher Scientific) was used for the normalization. All amplifications were carried out on StepOne Real-Time PCR System (Thermo Fisher Scientific). The changes in expression level between individual samples were determined by the ΔΔC_t_ method.

### Northern blot

Total RNAs isolated by TriPure reagent were separated along with Millennium RNA marker (Thermo Fisher Scientific) in formaldehyde-containing 1% agarose gel in 1 × MOPS buffer. After transfer, the membranes were hybridized overnight with antisense oligo probes listed in the **S6 Table** end-labeled with T4-PNK (New England Biolabs) using ^32^P-γ-ATP (Perkin Elmer). Washed membranes were exposed to a phosphor imaging screen and scanned by GE Typhoon TRIO Imager (GE Healthcare). Obtained images were processed by Image Lab software (BioRad).

### Reporter plasmid construction

To generate minigene system for monitoring ORF70/K3 RNA splicing and protein expression, the ORF70/K3 locus was amplified with oVM454 and oVM455 primers (see **S6 Table** for details) on BCBL-1 total DNA allowing its in-frame cloning into pCMV-FLAG-myc-22 (Sigma-Aldrich) using *Kpn*I and *Not*I restriction sites.

### Western blot

Total proteins, prepared by cells direct lysis in SDS protein sample buffer, were separated in 4-12% Bis-TRIS NuPAGE gels (Thermo Fisher Scientific) in 1 × MES buffer and transferred onto a nitrocellulose membrane. The in-house produced anti-ORF57 polyclonal antibody was used to ORF57 expression [68]. The anti-ORF50 antibody was generously provided by Dr. Yoshihiro Izumiya (University of California, Davis). The anti-Cas9 mouse monoclonal antibody was purchased from Cell Signaling (cat. no. 14697). The anti-FLAG polyclonal antibody (F7421, Sigma-Aldrich) was used for FLAG-ORF70 and anti-c-myc clone 9E10 antibody (a generous gift from Dr. Xuefeng Liu, Georgetown University) for K3-myc fusion detection in the reporter assay. The cellular β-tubulin detected by mouse monoclonal antibody (T5201, Sigma-Aldrich) was used as a loading control. The secondary horseradish peroxidase-labeled secondary antibodies (Sigma-Aldrich) were used for signal development using ProSignal Pico ECL Reagent (GeneSee Scientific) captured by Chemidoc Touch Imager (BioRad).

### Prediction and detection of circRNAs

KSHV circRNAs were predicted by CIRI2 pipeline (https://sourceforge.net/projects/ciri/) [37, 38] with default setting using RNA-seq reads from BCBL-1 WT B4 and 57KO #6 cells without or with valproic acid treatment. To experimentally confirm viral circRNAs expression, two µg of total RNA were treated with 5 U of RNase R (cat. # RNR07250, Lucigen, Middleton, WI) for 40 min at 37 °C. The reactions containing the corresponding amount of RNA without RNase R treatment served as a negative control. The obtained RNAs were reverse transcribed with SuperScript IV (cat. # 18091050, Thermo Fisher Scientific) at 50 °C for 1 h in the presence of random hexamers followed by detection of circular RNA as described [69]. For circRNA detection, 500 ng of cDNA were amplified by 40 PCR cycles using an inverse primer pair SuperFi II DNA polymerase (Thermo Fisher Scientific). The linear RNAs were amplified using 100 ng of cDNA for 25 cycles of PCR amplification using a corresponding primer pair. All primers are listed in the **S6 Table**.

## Supporting information

**S1 Fig.**
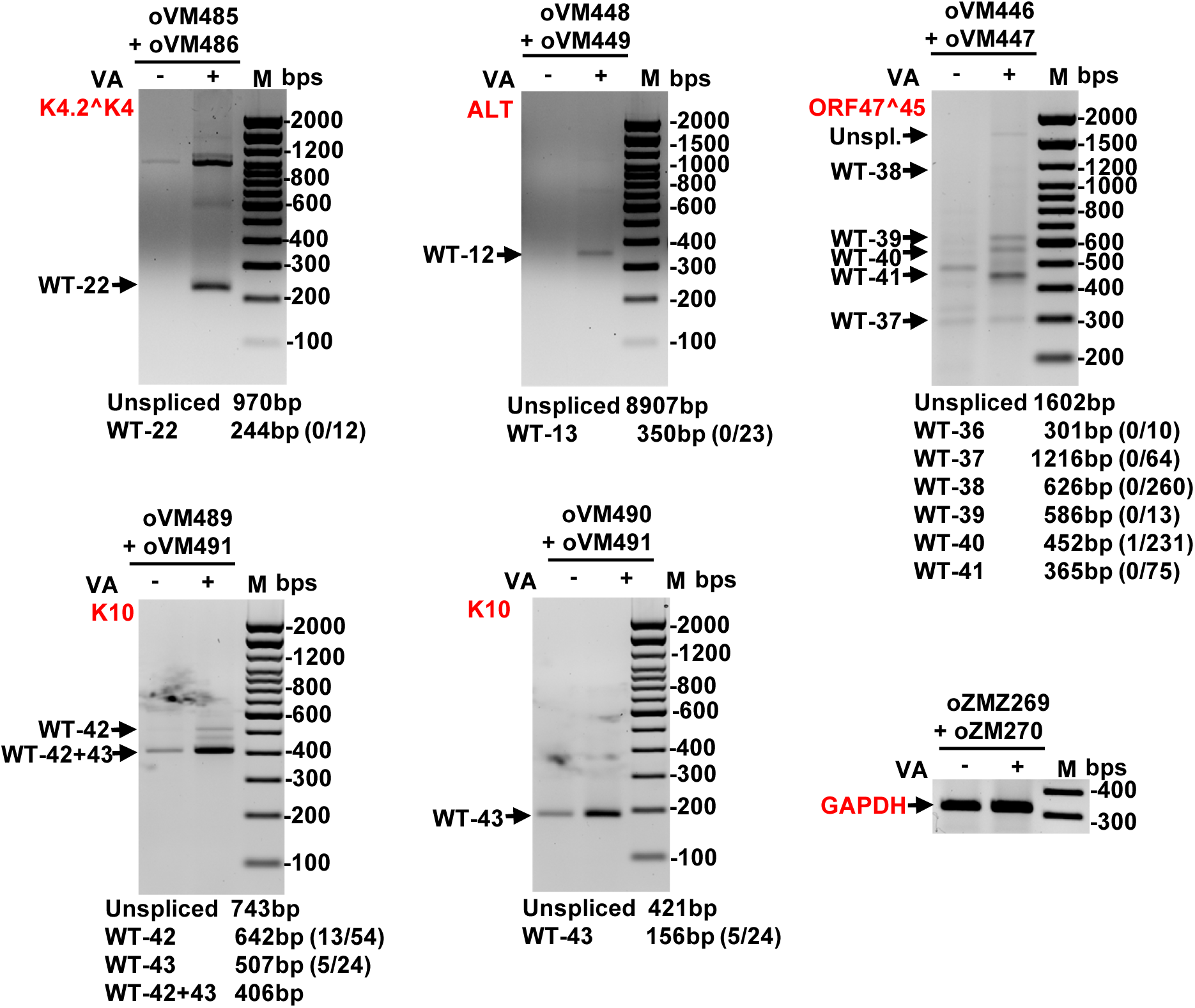
Selective validation of the newly identified splicing junctions in BCBL-1 WT cell. Total RNA from BCBL-1 cells without (VA-) or with (VA+) virus lytic replication was used as an RT-PCR template. For primer’s information, see Fig 3 **and S6 Table**.

**S2 Fig.**
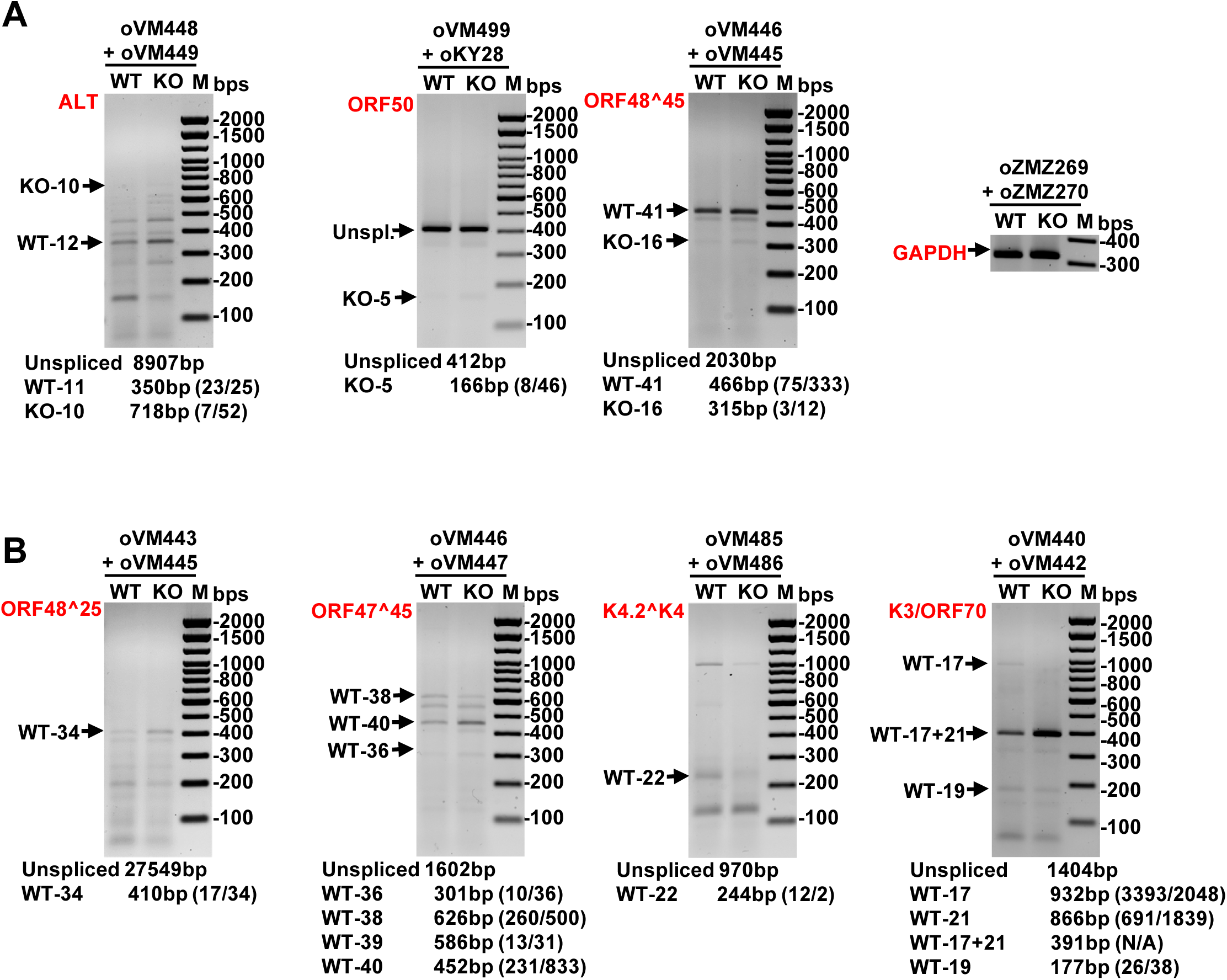
RT-PCR validation of selected RNA splicing events from the 57KO genome in BCBL-1 cells. Total RNA from WT and 57KO cells treated with valproic acid (VA) for 24 h to induce KSHV lytic replication was used in RT-PCR to detect 57KO-specific (KO) RNA splicing events (A) or splicing events with altered splicing efficiency between WT and 57KO genomes (B). For primer’s information, see Fig 3 **and S6 Table**.

**S3 Fig.**
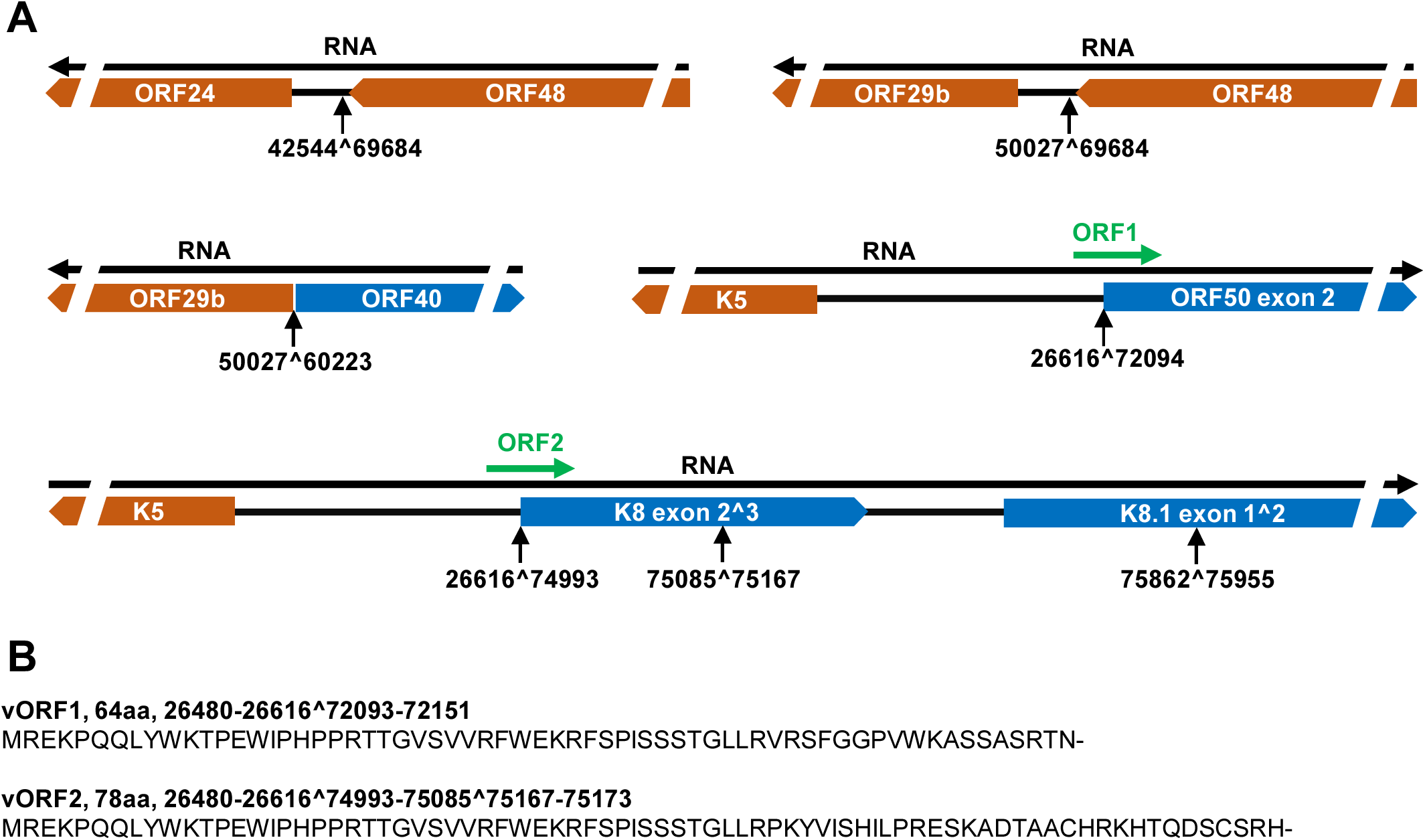
Viral long-range RNA splicing creates possible novel ORFs for KSHV coding potentials. (A) Novel transcripts (black arrows) generated by long-range RNA splicing identified in KSHV lytic infection. The annotated KSHV ORFs are shown in orange and blue. The novel ORFs (green arrows) spanning the long-range splicing junctions shown in Fig 6B were predicted using an ORF prediction program (ORFfinder, NCBI) with an AUG initiation codon ORF encoding at least 50 aa or more from all three frames. (B) The nucleotide position and amino acid sequences of the newly identified ORFs.

**S4 Fig.**
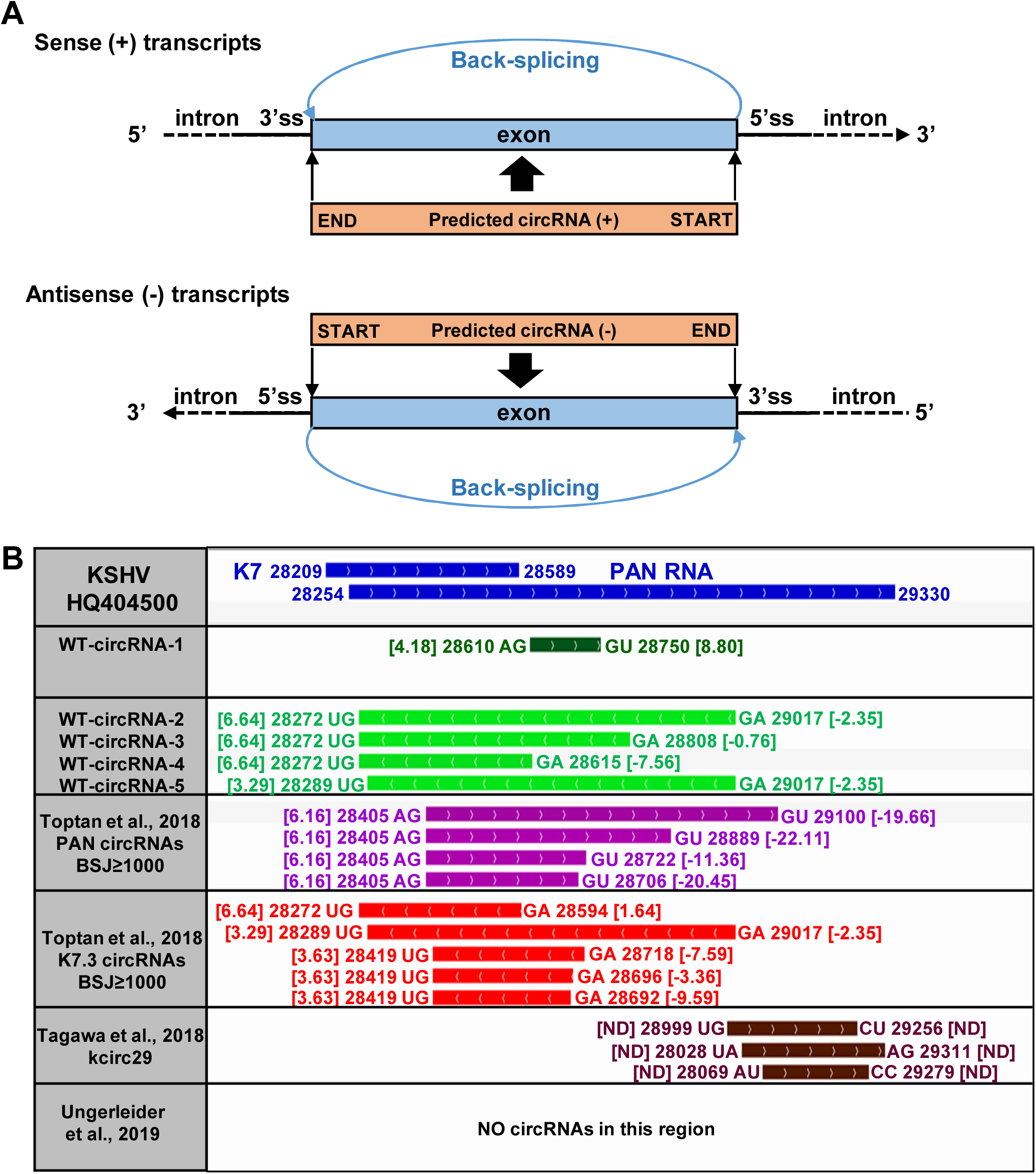
Lack of confident RNA splice sites in the PAN region to support predicted PAN-derived circRNAs. (A) A diagram of predicted circRNAs from the sense (+) or antisense (-) RNA transcripts expressed from the KSHV genome and their relations to the corresponding splice sites (ss). (B) The KSHV K7/PAN RNA locus in Bac36/BCBL-1 reference genome HQ404500 with the position and orientation of each circRNA predicted in this study (WT- circRNA-1-5) and other three reports ([34] supported with ≥1000 backsplicing junction reads (BSJ), [35]kcirc29, and [36]). The nucleotide position of each RNA splice site contributing to predicted circRNA biogenesis is shown together with the flanking dinucleotides and its splice site strength score (shown in parentheses) based on MAX:ENT model [31]. ND-not determined.

**S1 Table** Differential expression of KSHV genes in BCBL-1 single-cell clones carrying a WT B4 or 57KO #6 genome during latent and lytic infection determined by Limma Voom package based on the number of mapped RNA-seq reads in each group (A, B, or C) as described in Fig 1A.

**S2 Table** The summary table of all viral splicing junctions detected in individual groups (A, B, or C) described in Fig 1A.

**S3 Table** The prediction of KSHV circRNA in individual samples from WT and 57KO BCBL-1 upon lytic replication detected by CIRI2 pipeline in individual groups (A, B, or C) described in Fig 1A.

**S4 Table** Strength score analysis of the splice sites for linear RNA splicing junctions with at least two splicing junction reads mapped to the PAN RNA region from either plus- or minus-strand of KSHV genome. The splice site strength was determined by MAX:ENT algorithm [31].

**S5 Table** Strength score analysis of the splice sites flanking the predicted circRNAs from PAN and K7.3 RNAs with at least 1000 backsplicing junction reads from [34]. The splice sites strength was determined by MAXENT algorithm [31].

**S6 Table** The sequence and positions of oligos used in the study.

## Author Contributions

**Conceptualization:** Vladimir Majerciak, Zhi-Ming Zheng.

**Data curation:** Vladimir Majerciak, Alexei Lobanov, Maggie Cam.

**Formal analysis:** Vladimir Majerciak, Alexei Lobanov, Zhi-Ming Zheng.

**Funding acquisition:** Zhi-Ming Zheng.

**Investigation:** Vladimir Majerciak, Zhi-Ming Zheng, Alexei Lobanov.

**Methodology:** Vladimir Majerciak, Alexei Lobanov.

**Visualization:** Vladimir Majerciak.

**Writing – original draft:** Vladimir Majerciak, Zhi-Ming Zheng.

**Writing – review & editing:** Vladimir Majerciak, Zhi-Ming Zheng, Alexei Lobanov, Maggie Cam.

